# Parallel control of mechanosensory hair cell orientation by the PCP and Wnt pathways

**DOI:** 10.1101/527937

**Authors:** Joaquin Navajas Acedo, Matthew G. Voas, Richard Alexander, Thomas Woolley, Jay R. Unruh, Hua Li, Cecilia Moens, Tatjana Piotrowski

## Abstract

Cell polarity plays a crucial role during development of vertebrates and invertebrates. Planar Cell Polarity (PCP) is defined as the coordinated polarity of cells within a tissue axis and is essential for processes such as gastrulation, neural tube closure or hearing. Wnt ligands can be instructive or permissive during PCP-dependent processes, and Wnt pathway mutants are often classified as PCP mutants due to the complexity and the similarities between their phenotypes. Our studies of the zebrafish sensory lateral line reveal that disruptions of the PCP and Wnt pathways have differential effects on hair cell orientations. While mutations in PCP genes cause random orientations of hair cells, mutations in Wnt pathway members induce hair cells to adopt a concentric pattern. We show that PCP signaling is normal in hair cells of Wnt pathway mutants and that the concentric hair cell phenotype is due to altered organization of the surrounding support cells. Thus, the PCP and Wnt pathways work in parallel, as separate pathways to establish proper hair cell orientation. Our data suggest that coordinated support cell organization is established during the formation of lateral line primordia, much earlier than the appearance of hair cells. Together, these finding reveal that hair cell orientation defects are not solely explained by defects in PCP signaling and that some hair cell phenotypes warrant reevaluation.

## INTRODUCTION

Cell polarity is crucial for the function of many tissues and its establishment during development and morphogenesis has fascinated many generations of biologists. In addition to the well-studied apico-basal polarity of cells, cells are also coordinately aligned in the plane of a tissue’s axis, termed Planar Cell Polarity (PCP). PCP relies on the asymmetric distribution of ‘core PCP’ components, such as Van Gogh/Vangl, Frizzled, Dishevelled, Prickle/Spiny-Legs, Fmi/Celsr and Diego (Butler and Wallingford, 2017; Devenport, 2014; Goodrich and Strutt, 2011; Strutt and Strutt, 2005). This pathway was discovered in insects (Gubb and Garcia-Bellido, 1982; Lawrence, 1966; Lawrence and Shelton, 1975; Wong and Adler, 1993) and subsequently also identified in vertebrates (Heisenberg et al., 2000; Tada and Smith, 2000; Wallingford et al., 2000). Over twenty years of work have shown that PCP is required during key developmental processes that shape the embryo, such as gastrulation and neural tube closure (Butler and Wallingford, 2017; Devenport, 2014; Goodrich and Strutt, 2011). Despite the substantial progress that has been made in both invertebrates and vertebrates, how PCP is initiated is poorly understood. One proposed mechanism is that Wnt gradients, which can act as instructive morphogens in invertebrates and vertebrates, initiate the molecular asymmetry of PCP components (Chu and Sokol, 2016; Gao, 2012; Gros et al., 2009; Humphries and Mlodzik, 2018; Wu et al., 2013; Yang and Mlodzik, 2015). However, in some contexts Wnt ligands are required but not instructive (permissive) to drive PCP-dependent processes (Gordon et al., 2012; Ulrich et al., 2005; Witzel et al., 2006). In this study we set out to investigate the role of Wnt signaling in polarizing sensory hair cells.

The vertebrate inner ear is a classical model to study the function of the PCP pathway, as stereociliary bundle coordination depends on, and is very sensitive to changes in PCP (Deans, 2013; May-Simera and Kelley, 2012). Wnt gradients have been implicated in establishing PCP in the ear (Dabdoub et al., 2003; Qian et al., 2007) but due to the inaccessibility of the ear, the study of the function of the PCP and Wnt signaling pathways in coordinating hair cell alignment is challenging to investigate.

A more experimentally accessible model to study hair cell orientation is the sensory lateral line system of aquatic vertebrates that detects water movements across the body of the animal (Ghysen and Dambly-Chaudiere, 2007; Pujol-Marti and Lopez-Schier, 2013). Because of its superficial location in the skin, the lateral line is amenable to experimental manipulations and live imaging (Venero Galanternik et al., 2016). The lateral line system consists of volcano-shaped sensory organs (neuromasts) that are composed of mantle cells on the outside and support cells and mechanosensory hair cells in the center (Fig. 1a, b; (Ghysen and Dambly-Chaudiere, 2007)). The mechanosensory hair cells are homologous to the ones found in the inner ear (Chagnaud et al., 2017; Fritzsch and Straka, 2014). The sensory organs are derived from several migratory neurogenic, cephalic placodes/primordia that either migrate into the trunk, in the vicinity of the ear or into the head (reviewed in (Piotrowski and Baker, 2014)). As they migrate, primordia periodically deposit clusters of cells that later differentiate into sensory organs (Gompel et al., 2001; Ledent, 2002; Metcalfe et al., 1985). PrimordiumI (primI) and primordiumII (primII) both migrate into the trunk but arise from different placodes (primary placode and D0 placode, respectively). The D0 placode also gives rise to a third primordium that migrates onto the dorsal side of the trunk, called primD (Ghysen and Dambly-Chaudiere, 2007; Nunez et al., 2009).

**Figure 1.**
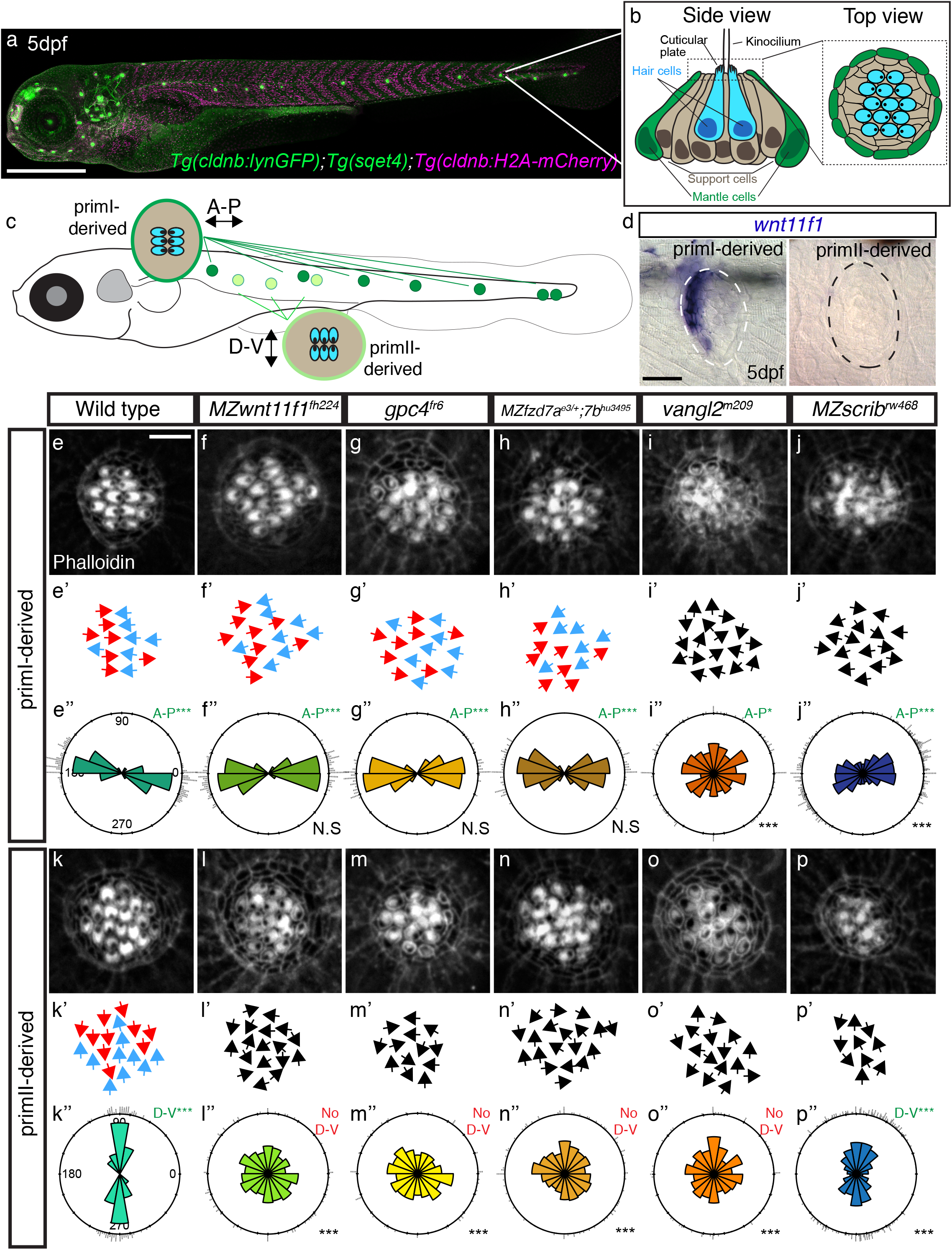
Mutations in the Wnt and PCP pathways affect the coordinated hair cell alignment in the lateral line. a. Confocal image of a 5 days post fertilization (dpf) zebrafish larvae transgenic for *Tg(cldnb: lynGFP), Tg(sqet4)* and *Tg(cldnb:H2A-mCherry)*. b. Diagram of the lateral view of a 5dpf neuromast showing the different cell types. c. Diagram of a 5dpf larva showing the two different orientations (A-P and D-V) of the hair cells in primI and primII-derived neuromasts, respectively. d. *In situ* hybridization of *wnt11f1* mRNA on primI and primII-derived neuromasts at 5dpf in wild type fish. e-j. Phalloidin staining showing hair cell orientation in primI-derived neuromasts of wild type (e), Wnt pathway mutants (f-h) and PCP mutants (i-j; p-val for *vangl2* primI= 7.33e-28, p-val for *MZscrib* primI= 1.41e-17). e’-j’. Depiction of individual hair cell orientation in each of the conditions tested in e-j. ln red are shown hair cells pointing posteriorly and in blue the ones that point anteriorly. Black arrows denote disruption of the wild type orientation (see e’’-j’’). e’’-j’’. Rose diagrams showing the hair cell orientation distribution respect to the longitudinal axis of the animal (horizontal) (WT n=194 hair cells, *MZwnt11f1* n=353, *gpc4* n=222, *MZfzd7a/7b* n=74, *vangl2* n=226, *MZscrib* n=535). On the bottom right is shown the Fisher’s exact test comparison with respect to the wild type for each condition. On the top right is shown the binomial test for each condition. k-p. Phalloidin staining showing hair cell orientation in primII-derived neuromasts of wild type (k), Wnt pathway mutants (l-n; p-val for *MZwnt11f1* primII= 1.69e-23, p-val for *gpc4* primII= 6.93e-31, p-val for *MZfzd7a/7b* primII= 3.91e-33) and PCP mutants (o-p, p-val for *vangl2* primII =2.52e-21, p-val for *MZscrib* primII= 9.22e-12). k’-p’. Depiction of hair cell orientation in k-p. Color code is the same as in e’-j’. k’’-p’’. Rose diagrams showing the hair cell orientation distribution with respect to the longitudinal axis of the animal (horizontal)(WT n=209 hair cells, *MZwnt11f1* n=295, *gpc4* n=214, *MZfzd7a/7b* n=182, *vangl2* n=141, *MZscrib* n=392). For e’-j’ comparisons of angle distribution with respect to wild type, ***=p<0.001, N.S= Not significative; A-P***= p<0.001, A-P*=p<0.05. e’’-p’’. A-P= Anteroposterior orientation. D-V= Dorsoventral orientation. Scale bar in a=500μm, d=20μm, e= 5μm.

Hair cells possess short actin-rich stereocilia adjacent to a tubulin-rich, long kinocilium. Each hair cell is planar polarized with the long microtubule-based kinocilium localized to one pole of the cuticular plate (Fig. 1b). Within a sensory organ, hair cells arise in pairs with their kinocilia pointing toward each other but aligned along a common axis (Fig. 1c). The positioning of the kinocilium along this axis is controlled by Notch signaling/Emx2 and its loss causes all hair cells within a neuromast to point into the same direction (Jacobo A, 2018; Jiang et al., 2017). Hair cells within sensory organs show different polarities of this common axis depending on which placode/primordium they originate from, allowing the animal to sense water motion coming from all directions (Fig. 1c, (Lopez-Schier et al., 2004)). In primI-derived trunk neuromasts, hair cells are planar polarized along the anterioposterior (A-P) axis and hair cells in primII-derived neuromasts are aligned along the dorsoventral (D-V) axis. The differential hair cell orientation has been correlated with the direction of migration of the different primordia from which these sensory organs originate from, however no underlying mechanisms have been identified (Ghysen and Dambly-Chaudiere, 2007; Lopez-Schier et al., 2004; Nunez et al., 2009).

At the single cell level, previous reports have suggested that PCP regulates hair cell orientation in the pLL of zebrafish by controlling cell division angles and cell rearrangements of the progenitors that form the hair cells (Lopez-Schier and Hudspeth, 2006; Mirkovic et al., 2012). Yet the molecular code for how different neuromast axial orientations and individual hair cell orientations are coordinated remains ill understood.

Here, we show that loss of PCP and Wnt pathway genes have different consequences on hair cell orientations. While mutations in the PCP signaling genes *vangl2* and *scrib* cause disorganized hair cell orientations in all neuromasts, mutations in the Wnt pathway genes *wnt11f1* (formerly known as *wnt11r), gpc4* and *fzd7a/7b* show a striking concentric pattern of hair cell orientation in a subset of neuromasts. As neither the core PCP component Vangl2, nor Notch/Emx2 signaling are affected in Wnt pathway mutants we conclude that the Wnt pathway acts in parallel to these pathways. In addition, the concentric hair cell phenotype in Wnt pathway mutants is caused by the disruption of coordinated organization of the surrounding support cells, rather than by affecting the axis of poloarity or kinocilium positioning in individual hair cells.

Interestingly, the expression patterns of Wnt pathway genes in mature neuromasts suggests that the Wnt pathway acts very early in lateral line development. Thus, Wnt signaling does not instruct PCP, but rather acts to coordinate support cell organization during the formation and migration of the primordium before the appearance of hair cells. Overall, our findings demonstrate that hair cell orientation defects cannot solely be attributed to defects in the PCP pathway and that some phenotypes formerly characterized as PCP defects need to be re-evaluated.

## RESULTS

### Mutations in Wnt and PCP pathway genes cause different hair cell orientation phenotypes

We chose to study how primI and primII-derived neuromast orientations are established because during a large *in situ* screen, we unexpectedly found asymmetric distributions of the Wnt ligand *wnt11f1* (formerly known as *wnt11r, wnt11-related*, Postlethwait et al., in preparation), the ortholog of mammalian WNT11. *wnt11f1* is expressed in cells along the anterior edge of only primI-derived neuromasts, but is absent from primII-derived neuromasts (Fig. 1d, Fig. Suppl. 1).

Since Wnt ligands can instruct planar polarization of cells and hair cells (Bradley and Drissi, 2011; Chu and Sokol, 2016; Gao et al., 2011; Gros et al., 2009; Wada and Okamoto, 2009a, b; Wallingford et al., 2000), we hypothesized that *wnt11f1* establishes hair cell orientation by directing PCP in primI-derived neuromasts. We measured hair cell orientation in the cuticular plate using Phalloidin staining, which labels the actin-rich stereocilia but not the tubulin-rich kinocilium (Fig. 1b). We used the position of the kinocilium to determine the axis of polarity of each hair cell. Phalloidin stainings of sibling primI-derived neuromasts show that hair cells have a significant bias in orientation parallel to the A-P axis (Fig. 1e-e’) based on the the angles with respect to the horizontal in rose diagrams (Fig. 1e’’). In contrast, primII-derived neuromasts show a strong orientation bias along the D-V axis (Fig. 1k-k’’). Furthermore, neighboring hair cells in both primordia show coordinated polarities (Fig. Suppl 2b, h). Surprisingly, zygotic and maternal zygotic mutations in *wnt11f1* do not affect hair cell orientation in primI-derived neuromasts in which *wnt11f1* is expressed (Fig1. f-f’, p=1; Fig. Suppl. 2c), but disrupt the wild type hair cell orientation in primII-derived neuromasts (Fig1. l-l’’; Fig. Suppl. 2i).

*gpc4* (*glypican4*, a heparan sulfate proteoglycan (HSPG)) and *fzd7* (a frizzled-class receptor) interact with Wnt ligands during convergent extension (CE) and we tested whether mutations in these genes also cause hair cell polarity defects (Djiane et al., 2000; Ohkawara et al., 2003; Rochard et al., 2016; Roszko et al., 2015; Topczewski et al., 2001; Witzel et al., 2006). Indeed, *gpc4* and maternal zygotic (MZ) *MZfzd7a/7b* mutants show the same concentric hair cell phenotype as *MZwnt11f1* mutants suggesting they act in the same pathway (Fig. 1g-g’’, m-m’’; and h-h’’, n-n’’; Fig. Suppl. 3b-f). This interpretation is supported by a double mutant analysis between *wnt11f1* and *gpc4*. Double *wnt11f1;gpc4* homozygous larvae do not show an additive phenotype and only possess hair cell defects in primII-derived neuromasts (Fig. Suppl. 2n). We therefore refer to *wnt11f1, gpc4* and *fzd7a/7b* as ‘Wnt pathway genes’. *gpc4* and *fzd7a/7b* interact with *wnt11f2* (formerly called *wnt11/silberblick*, Postlethwait et al., in prep.) in convergent extension (Capek et al., 2019; Petridou et al., 2018; Witzel et al., 2006), we therefore wondered if *wnt11f2* mutants also possess hair cell defects and if *wnt11f2* possibly interacts with *wnt11f1*. However, *wnt11f2* mutants have normal hair cell orientations, as do *wnt11f1;wnt11f2* double heterozygous animals, indicating that these two paralogs do not interact (Fig. Suppl. 2o).

Since *gpc4* and *MZfzd7a/7b* mutants have been described as PCP signaling mutants, we compared the phenotypes of *MZwnt11f1, gpc4* and *MZfz7a/7b* mutants to the hair cell phenotype of the PCP mutants *vangl2* and *scribble (scrib)*. It was previously shown that a mutation in the core PCP gene *vangl2* disrupts the hair cell orientation in both primI and primII-derived neuromasts (Fig. 1i-i’’, o-o’’, (Lopez-Schier and Hudspeth, 2006; Mirkovic et al., 2012); Fig. Suppl. 2f, l). We also observed that hair cells in maternal-zygotic mutants for *scrib (MZscrib)* show a significant deviation of hair cell orientations in both primI and primII-derived neuromasts (Fig. 1j-j’’, p-p’’). However, the phenotype is not as severe as in *vangl2* mutants and primI and primII neuromasts still show a bimodal distribution along the A-P and D-V axes, respectively (Fig. 1j”, p”; Fig. Suppl. 2g, m). *scrib* likely acts partially redundantly with other PCP proteins. These results suggest that while PCP genes are required to establish proper hair cell orientation in all neuromasts, Wnt pathway genes act to establish hair cell orientation only in primII-derived neuromasts.

Mutations in both Wnt and PCP pathway genes result in misorientation of hair cells. However, by measuring the hair cell orientation defects in Wnt pathway and PCP signaling mutants we found that mutations in Wnt pathway genes cause hair cells in primII-derived neuromasts to orient in a striking concentric fashion (Fig. 2b, g, Fig. Suppl. 3b-f), whereas in *vangl2* mutants hair cells are oriented randomly (Fig. 2c-c’, h-h’, Fig. Suppl. 2f, l). Likewise, hair cells in the other PCP signaling mutant *MZscrib* do not arrange in a concentric fashion (Fig. Suppl. 3d, g). This difference in phenotype suggests that the Wnt and PCP signaling pathways control hair cell orientation in parallel, rather than through a common pathway. However, Wnt and PCP pathways have been described to interact genetically in the establishment of hair cell orientation in the inner ear (Qian et al., 2007). To assess a possible genetic interaction between *vangl2* and *wnt11f1* in lateral line neuromasts we generated double homozygous *vangl2* and *MZwnt11f1* larvae. In double homozygous fish, primI-and II-derived neuromasts show random hair cell orientation (Fig. 2e, j; Fig. Suppl. 4b, d). In addition, the concentric phenotype shown by primII neuromasts of *MZwnt11f1* single mutants (Fig. 2i, Fig. Suppl. 4c) disappears in the double *MZwnt11f1;vangl2* mutants (Fig. 2j; Fig. Suppl. 4d). Thus, the disorganized hair cell phenotype caused by a mutation in *vangl2* is epistatic over the concentric phenotype, providing additional support for the idea that the PCP and Wnt pathways may work in parallel and have different roles during the establishment of hair cell orientation in primII-derived neuromasts.

**Figure 2.**
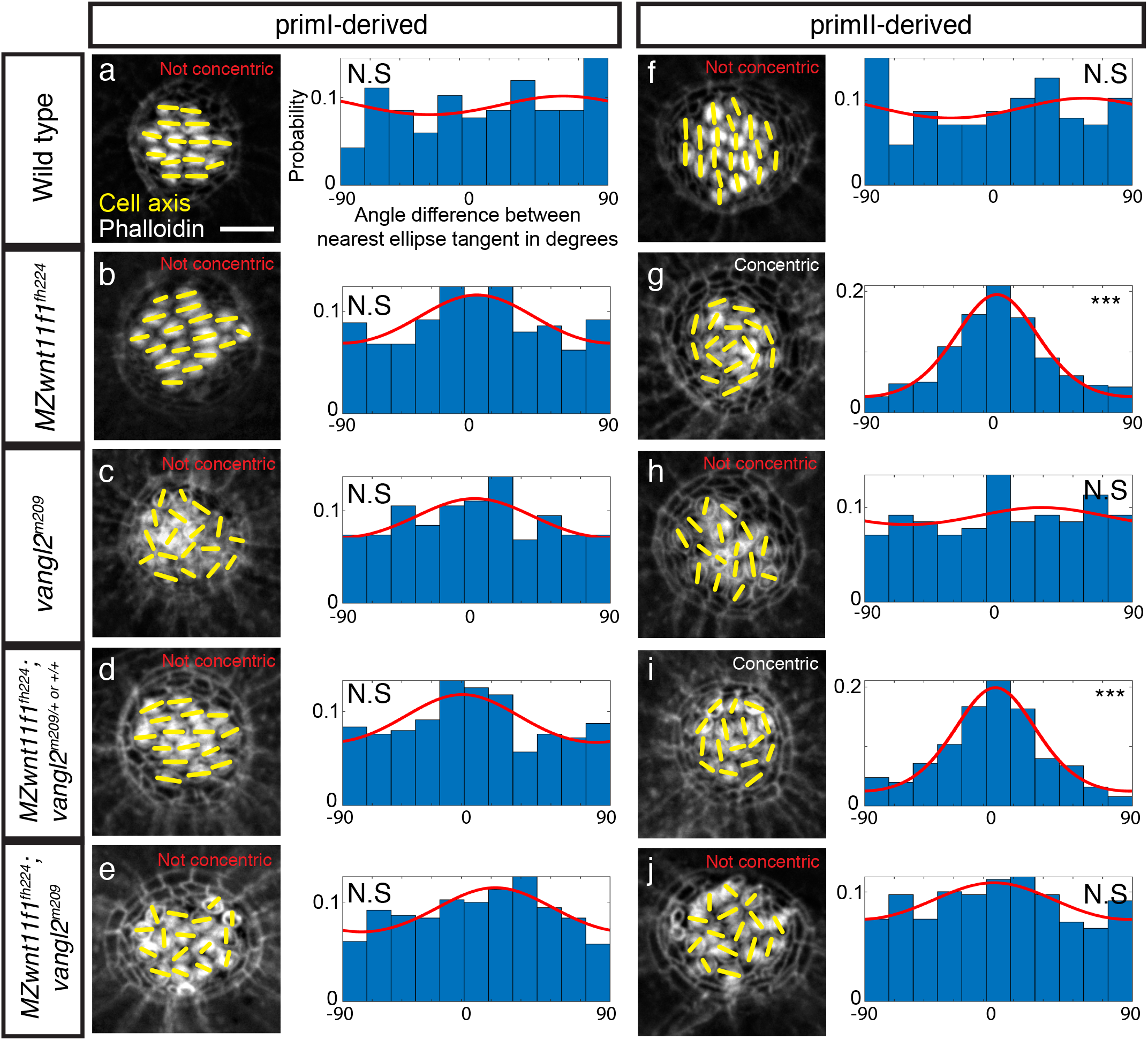
Hair cells show a disorganized pattern in PCP mutants, while they show a concentric orientation pattern in primII of Wnt pathway mutants. a-e. Phalloidin images showing the cell polarity axis (yellow lines) in primI-derived neuromasts of wild type (a), Wnt pathway mutant *MZwnt11f1* (b), PCP mutant *vangl2* (c), single *MZwnt11f1* siblings (d) and double *MZwnt11f1;vangl2* (e) homozygous mutants and their distribution of angles with respect to the nearest ellipse tangent (concentricity). Note that none of the conditions shows significative concentricity (Uniform distribution p-values in a= 0.423, b= 0.054, c= 0.464, d= 0.165, e= 0.077, f= 0.019; WT n=194 hair cells, *MZwnt11f1* n=353, *vangl2* n=226, *MZwnt11f1;vangl2* sibling n=276, *MZwnt11f1;vangl2* doubles n=406). The Von Mises distribution p-value and the R^2^ fit were calculated for each of the conditions but none show a good fit. f-j. Phalloidin images showing the cell polarity axis (yellow lines) in primII-derived neuromasts of wild type (f), Wnt pathway mutant *wnt11f1* (g), PCP mutant *vangl2* (h), single *MZwnt11f1* siblings (i) and double *MZwnt11f1;vangl2* (j) homozygous mutants and their distribution of angles with respect to the nearest ellipse tangent (concentricity). Note that only the Wnt pathway mutants (g, i) show significative concentricity (Uniform distribution p-values in f= 0.314, g= 1.28e-27, h= 0.86, i= 0.488, j= 02.92e-19, k= 0.298; WT n=209 hair cells, *MZwnt11f1* n=295, *vangl2* n=141, *MZwnt11f1;vangl2* sibling n=252, *MZwnt11f1;vangl2* doubles n=365). The Von Mises distribution p-value and the R^2^ fit were calculated for each of the conditions, and only the Wnt pathway mutants show a good fit (R^2^ for g= 0.968, j= 0.961). Yellow lines in a-j indicate the hair cell polarity axis, determined by the position of the kinocilium. Not Concentric vs Concentric labels were based on calculations in a-j. Scale bar= 5μm.

### Asymmetric Vangl2 localization is not affected in the hair cells of Wnt pathway mutants

Since loss of *vangl2* disrupts the concentric phenotype, we hypothesized that PCP signaling might be correctly established in hair cells of the Wnt pathway mutants. To test whether one of the landmarks of PCP signaling, asymmetric distribution of Vangl2 (Devenport, 2014), is disrupted in hair cells of Wnt pathway mutants we used an anti-Vangl2 antibody to detect endogenous levels in 5dpf neuromasts (Fig. 3). In wild type fish, Vangl2 is enriched on the posterior side of hair cells in primI-derived neuromasts (Fig. 3a-a’) and on the ventral side of hair cells in primII-derived neuromasts (Fig. 3e-e’, i). Approximately half of the hair cells show Vangl2 signal in the same pole with the kinocilium (Fig. 3i, Fig. Suppl. 5), which agrees with previous reports using a fluorescent reporter (Mirkovic et al., 2012). While Vangl2 staining is not detected in *vangl2* mutants (Fig. Suppl. 6), *MZscrib* mutant fish show reduced signal and fewer hair cells with asymmetric Vangl2 in primI and primII-derived neuromasts (Fig. 3b-b’, f-f’, i) compared with their siblings (Fig. 3c-c’, g-g’, i). The reduction in signal agrees with previous reports in the mouse cochlea (Montcouquiol et al., 2006) and might explain why hair cell disorganization is not as dramatic as in *vangl2* mutants. Additionally, the correlation of the signal with the axis of polarity defined by the kinocilium is disrupted in hair cells that show asymmetric Vangl2 (Fig. Suppl. 5). This observation further indicates that PCP signaling is affected in hair cells of *MZscrib*. In contrast, *MZwnt11f1* mutants possess asymmetric distribution of Vangl2 in hair cells of both primI and primII-derived neuromasts showing that PCP signaling is not affected (Fig. 3d-d’, h-h’, i; Fig. Suppl. 5). To further test whether hair cells in *MZwnt11f1* mutants show asymmetric Vangl2 localization, we generated a hair cell specific promoter-driven GFP-Vangl2, *Tg(myo6b:GFP-XVangl2)*. We find that in wild type fish, based on the asymmetric enrichment of GFP fluorescence around the circumference of the cuticular plate at the base of the stereo- and kinocilia, mosaic expression of fluorescent Vangl2 is asymmetrically localized in hair cells of primI and primII-derived neuromasts (Fig. 3j-l, p). Likewise, in *MZwnt11f1* mutants, GFP-Vangl2 is also asymmetrically localized around the circumference of each hair cell in both primI and primII-derived neuromasts (Fig. 3m-p). These results indicate that in contrast to the PCP mutant *vangl2*, individual hair cell PCP is normal in Wnt pathway mutants. This finding suggests that the concentric arrangement of hair cells in Wnt pathway mutants is caused via different mechanism than by a defect in PCP.

**Figure 3.**
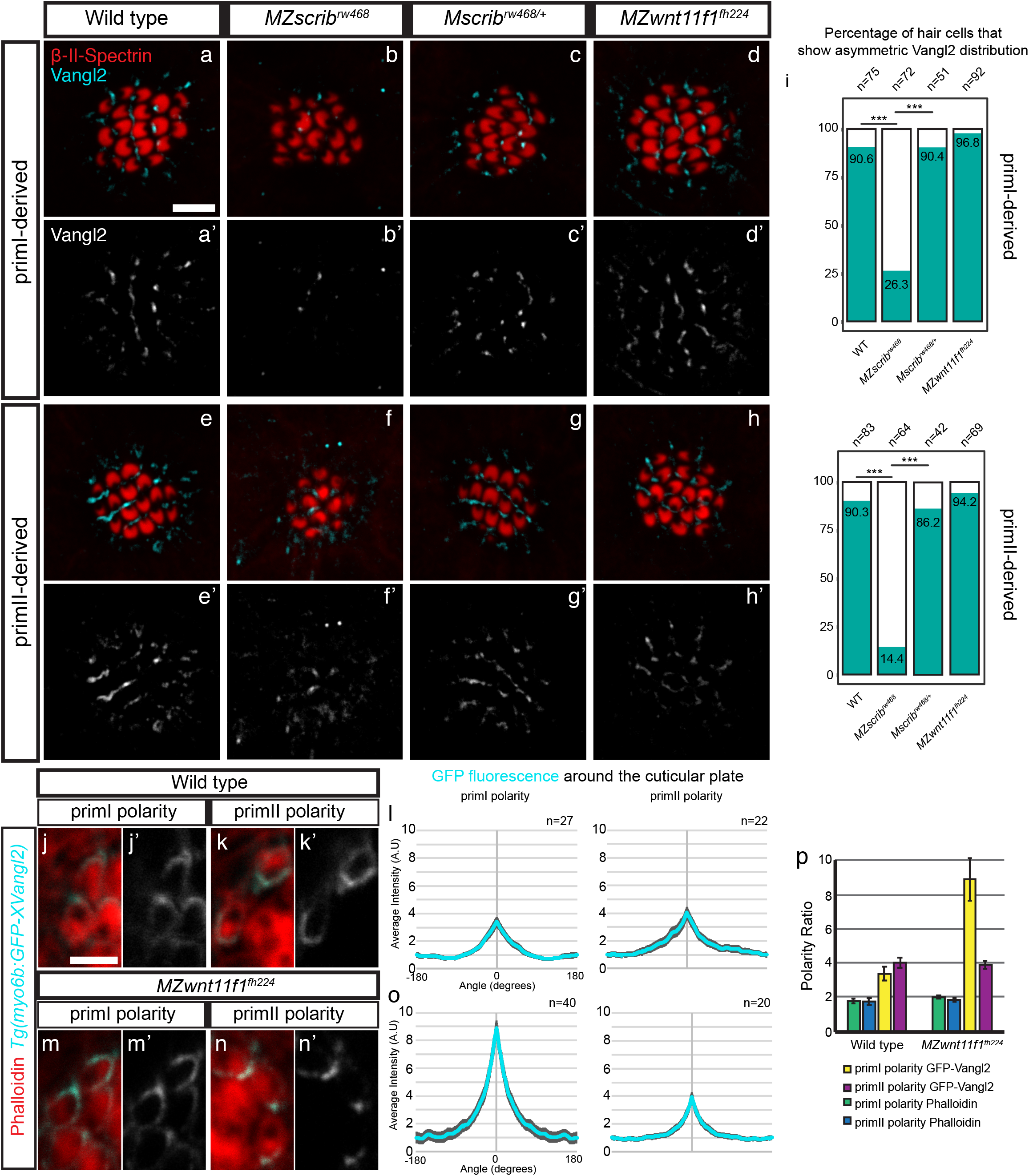
Vangl2 asymmetry is not affected in hair cells of *MZwnt11f1* mutants. a-d’. Double β-II-Spectrin (labeling the cuticular plates) and Vangl2 immunodetection in primI neuromasts of 5dpf wild type (a-a’), *MZscrib* mutants (b-b’), *Mscrib* siblings (c-c’) and *MZwnt11f1* mutants (d-d’). e-h’. Double β-II-Spectrin (labeling the cuticular plates) and Vangl2 immunodetection in primII neuromasts of 5dpf wild type (e-e’), *MZscrib* mutants (f-f’), *Mscrib* siblings (g-g’) and *MZwnt11f1* mutants (h-h’). i. Quantification of the percentage of hair cells that show asymmetric Vangl2 for each of the conditions in both primordia-derived neuromasts. j-n’. Clonal localization of GFP-XVangl2 in hair cells of neuromasts that show primI and primII polarity in wild type (j-j’, k-k’) and *MZwnt11f1* mutants (m-m’, n-n’) neuromasts. The majority of the clones were localized on neuromasts in the head, but these show the same phenotype as primI and primII-derived neuromasts in each of the cases (data not shown). l, o. Quantification of GFP-XVangl2 fluorescence around the cuticular plates of hair cells in neuromasts that show primII and primII polarity of wild type (l) and *MZwnt11f* mutants (o). p. GFP fluorescence and Phalloidin enrichment (Polarity Ratio) in the different conditions analyzed in (l, o). Error bars in l, o and p represent S.E.M. ***= Fisher’s exact test p<0.01. Scale bar in a= 5μm. Scale bar in j= 2μm.

### The phenotypes of Wnt and PCP pathway mutants arise at different times during development

The differences in the hair cell orientation phenotypes exhibited by Wnt and PCP pathway mutants suggest that they do not act in the same pathway. To identify further differences in their phenotypes we investigated PCP-dependent cell behaviors of the hair cell progenitors *in vivo*. The two daughters of a hair cell progenitor often change their position with respect to each other before differentiating (Lopez-Schier and Hudspeth, 2006; Mirkovic et al., 2012)(Suppl. Video 1). These cell rearrangements and their duration depend on functional PCP signaling during development and regeneration. In *vangl2* mutants the hair cell rearrangements do not lead to complete reversal of positions, they sometimes occur multiple times and their duration is prolonged (Mirkovic et al., 2012). In contrast, time-lapse analyses of the behavior of developing hair cell progenitors in primII-derived neuromasts of *MZwnt11f1* mutants reveals no significant differences in the number of times they rearrange, the duration of the rearrangements or the angle of divisions with respect to the radius of the neuromast compared to wild type (Fig. Suppl. 7c-e). Based on these results we conclude that hair cell pairs arise normally in *MZwnt11f1* mutants. We also identified differences in the timing of the onset of the hair cell phenotypes between PCP and Wnt pathway mutants. In wild type larvae a pair of hair cells develops from a common progenitor (Lopez-Schier and Hudspeth, 2006) with the kinocilia of both hair cells pointing towards each other in 96% of the cases (Fig. 4a). Interestingly, the first pair of hair cells in *MZwnt11f1* mutants shows correct orientation 69% of the time (Fig. 4b), while in *vangl2* mutants they orient properly only in 23% of the cases (Fig. 4c). Combined with the results from the cell rearrangement analyses we conclude that the majority of *MZwnt11f1* mutant hair cells have the proper orientation with respect to their sister hair cell as they form, whereas most *vangl2* mutant hair cells show defective hair cell polarities that are not coordinated with their sister hair cell as soon as the hair cell progenitor divides.

**Figure 4.**
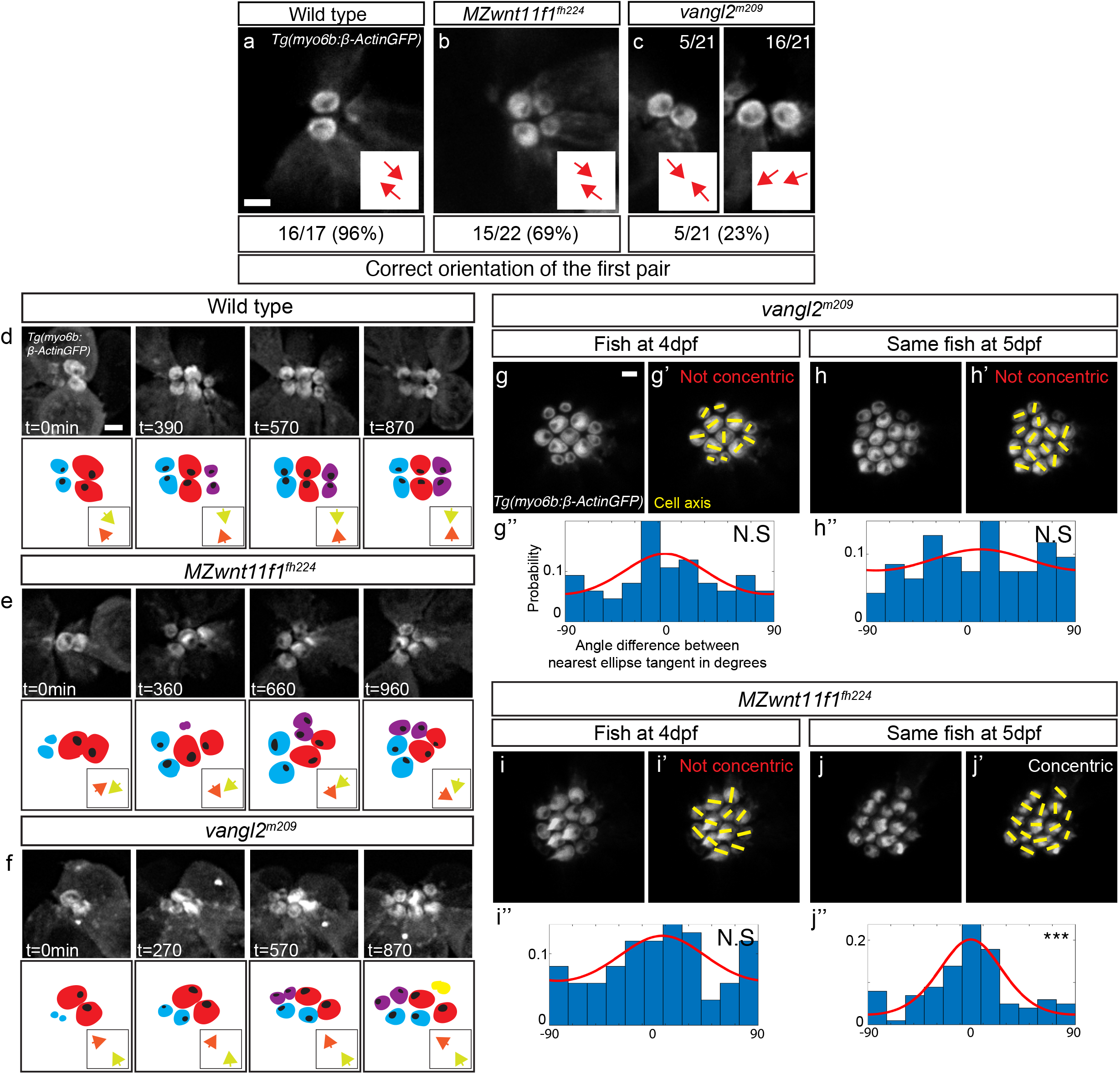
Differences in the appearance of the hair cell phenotypes in *vangl2* and *wnt11f1* mutants. a-c. Orientation of the first pair of hair cells in primII-derived neuromasts of wild type (a), *MZwnt11f1* mutants (b) and *vangl2* mutants (c) labeled with *Tg(myo6b:β-Actin-GFP)*. ln c, the left panel shows the number of correctly oriented pairs, and the right panel shows the number of incorrectly oriented pairs (the majority). The percentages for each of the conditions represent the number of pairs that show correct hair cell orientation. d-f. Hair cell orientation over time in wild type (n=3 animals) (d), *MZwnt11f1* mutants (n=5) (e) and *vangl2* mutants (n=2) (f) labeled with *Tg(myo6b:β-Actin-GFP)*. Each of the conditions is depicted using a color-coded cartoon, and the hair cell orientation of the first pair is represented by arrows in the box on the lower right. To differentiate between each hair cell of the pair, each of the arrows is in a different color. g-j’. Hair cell orientation of the same *vangl2* mutant fish at 4dpf (g-g’) and 5dpf (h-h’), and the same *MZwnt11f1* mutant embryo at 4dpf (i-i’) and 5dpf (j-j’) in a *Tg(myo6b:β-Actin-GFP)* transgenic background. g’’, h’’, i’’, j’’. Distribution of angles respect to the nearest ellipse tangent (concentricity) for primII-derived neuromasts of all the conditions tested. Note that only the *MZwnt11f1* mutant at 5dpf (j’, j’’) show significative concentricity (Uniform distribution p-values in g’’= 0.223, h’’= 0.469, i’’= 0.308, j’’= 1.97e-07). The Von Mises distribution p-value and the R^2^ fit were calculated for *MZwnt11f1* mutants at 5dpf, which shows a good fit (R^2^ for j’’= 0.749). Yellow lines represent the polarity axis for each hair cell. ‘Not Concentric’ vs ‘Concentric’ labels in g’-j’ were based on g’’, h’’, i’’ and j’’ (n=8 for each condition). Scale bars in a, d and g=2 μm

As *MZwnt11f1* mutant hair cells eventually all show orientation defects with respect to their sister hair cell, we asked if this phenotype arises progressively as sensory organs grow. We measured the polarity of the first hair cell pair over time as additional hair cells appear and observed that in wild type neuromasts the hair cells keep pointing towards each other or their polarity is even refined (Fig. 4d; Suppl. Video 2). In contrast, in *MZwnt11f1* mutants the correct orientation is progressively lost (Fig. 4e; Suppl. Video 3). In *vangl2* mutants, the aberrant polarity of the first pair also changes over time (Fig. 4f; Suppl. Video 4). At this point we do not understand if this effect of cell crowding on the Wnt and PCP phenoytpes is caused by the same mechanism or not.

As 5dpf *vangl2* hair cells do not arrange in concentric circles, we asked whether concentricity might be an intermediate state that is subsequently lost in PCP mutants. To test this possibility, we performed time lapse analyses of developing hair cells in *vangl2* and *MZwnt11f1* neuromasts between 4-5dpf. *vangl2* hair cells never arrange in a concentric fashion and are always randomly oriented (Fig. 4g-h’’, Fig Suppl. 8). Thus, concentricity is not a property of developing PCP-deficient hair cells at any stage of their development. Likewise, 4dpf *MZwnt11f1* (Fig. 4i-i’’) mutant hair cells are not organized in a concentric fashion, even though they are misaligned (Fig. Suppl. 8). However, starting at 5dpf, concentricity is detected in *MZwnt11f1* mutants (Fig. 4j-j’’, Fig. Suppl. 8). Therefore, concentricity arises over time in *MZwnt11f1* mutants as more hair cells are being added. We conclude that the hair cell defects in PCP and Wnt mutants are caused by different mechanisms, because the majority of *vangl2* mutant hair cells are already misaligned with respect to their sister hair cell after progenitor division, whereas the misalignment of the two sister hair cells in *MZwnt11f1* mutant hair cells arises over time.

### Wnt pathway genes are required to achieve coordinated support cell organization

As the hair cell defect arises over time in *MZwnt11f1* mutants we wondered if the hair cell phenotype could be secondary to defects in neighboring support cells. Support cells surround and sit underneath developing hair cells and serve as hair cell progenitors (Lopez-Schier and Hudspeth, 2006; Lush and Piotrowski, 2014; Romero-Carvajal et al., 2015). Interestingly, support cells are aligned along the A-P and D-V axes in primI and primII-derived neuromasts, respectively, raising the possibility that the Wnt pathway genes might be acting on them (Fig. 5a-b’).

**Figure 5.**
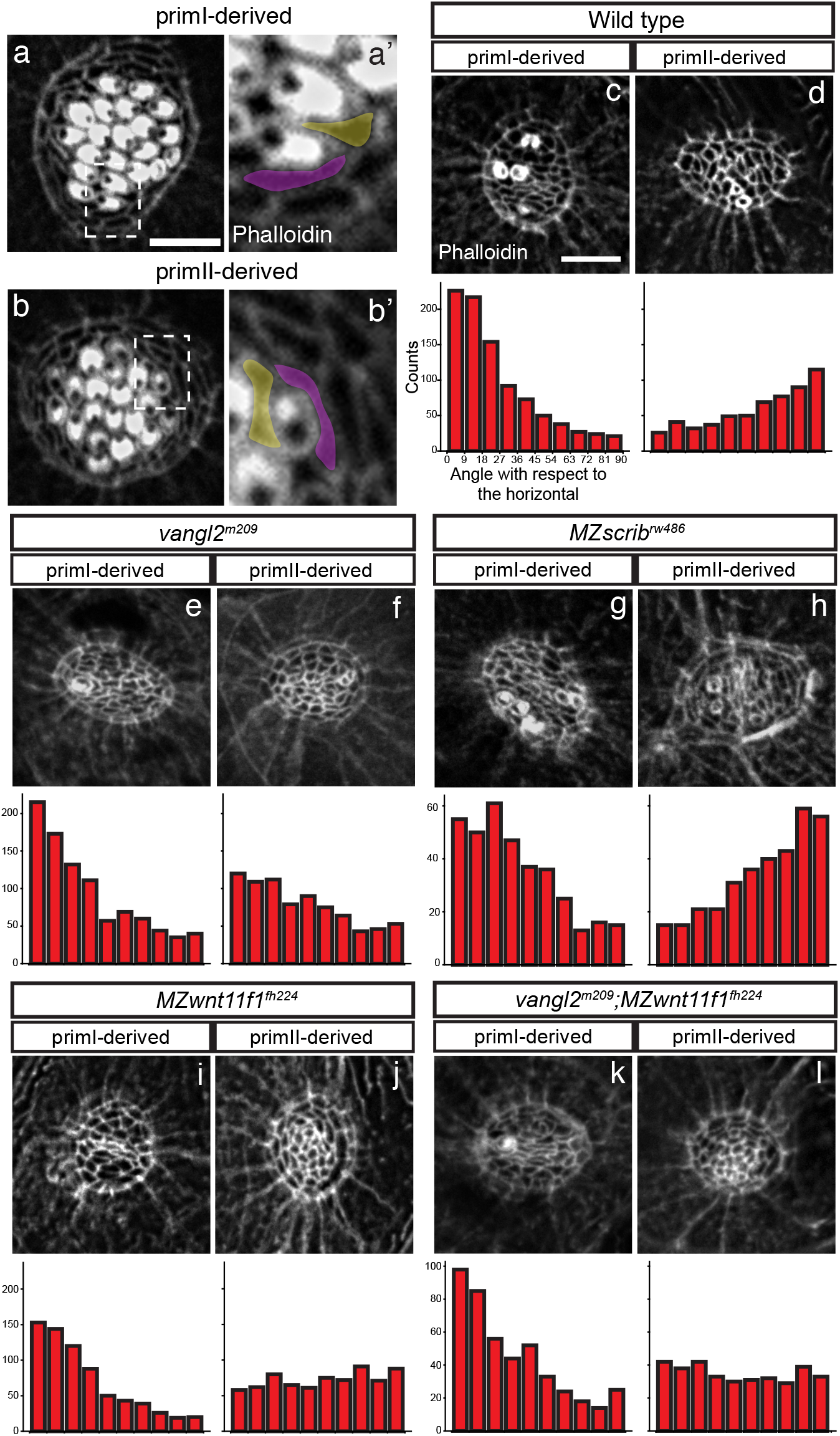
Wnt pathway genes are required for proper coordinated support cell organization. a-b’. Phalloidin staining of wild type primI and primII-derived neuromasts. The dotted square outlines the area magnified in a’ and b’. Two support cells each were false colored to depict their orientation. b-j. Phalloidin staining of neuromasts 3 hours after hair cell ablation. c-d. Wild type primI (c) and primII-derived (d) neuromasts. e-f. *vangl2* mutant primI (e) and primII-derived (f) neuromasts. g-h. *MZscrib* mutant primI (g) and primII-derived (h) neuromasts. i-j. *MZwnt11f1* mutant primI (i) and primII-derived (j) neuromasts. k-l. *vangl2;MZwnt11f1* double mutant primI (k) and primII-derived (l) neuromasts. The histograms show the distribution of binned angles of cell orientation with respect to the horizontal for each of the conditions tested. c-d. Angle distribution in Wild type primI (c, One-way chi square test p=1.78e-87) and primII-derived (d, p=4.54e-19) neuromasts. Note that primI distribution is skewed towards being horizontally aligned, while the distribution in primII-derived neuromasts is towards vertical alignment. e-f. Angle distribution in *vangl2* mutant primI (e, p=4.91e-13) and primII-derived (f, p=2.14e-16) neuromasts. g-h. Angle distribution in *MZscrib* mutant primI (g, p=1.40e-14) and primII-derived (h, p=9.60e-13) neuromasts. i-j. Support cell orientation in *MZwnt11f1* mutant primI (i, p=5.32e-52) and primII-derived (j, p=0.006) neuromasts. k-l. Support cell orientation in *vangl2;MZwnt11f1* double mutant primI (k, p=6.69e-26) and primII-derived (l, p=0.22) neuromasts. Scale bar in a= 5μm.

To better visualize any defects in support cell organization we removed hair cells by soaking the larvae in the antibiotic neomycin (Harris et al., 2003) and calculated their orientation with respect to the horizontal (Fig. Suppl. 9). In wild type larvae, support cells show a coordinated, elongated alignment along the horizontal plane (A-P axis) in primI-derived neuromasts (Fig. 5c-c’) and vertical plane (D-V axis) in primII-derived neuromasts (Fig. 5d-d’). These cell alignments demonstrate that hair cell ablation does not disrupt instrinsic support cell alignment. Unexpectedely, we observed that in *vangl2* mutants primI-derived support cells are normally aligned along the horizontal axis, even though their hair cell orientation is randomized (Fig. 5e-e’). *vangl2* mutant support cells in primII-derived neuromast still align, but horizontally (Fig. 5f-f’), a finding we currently cannot explain. However, the *MZscrib* PCP mutant shows wild type support cell organization in both primI (Fig. 5g-g’) and primII-derived neuromasts (Fig. 5h-h’). These results imply that functional PCP is not required for support cell alignment but that PCP genes act mostly in hair cells. On the other hand, in *MZwnt11f1* mutants, primI-derived support cells are horizontally aligned (Fig. 5i-i’), whereas support cells in primII-derived neuromasts do not possess an evident coordinated vertical organization (Fig. 5j-j’). To investigate a possible genetic interaction between PCP and Wnt pathway genes during the establishment of support cell organization, we asked whether the support cell phenotype shown by Wnt pathway mutants would disrupt the normal support cell alignment in PCP mutants. Double *vangl2;MZwnt11f1* homozygous mutants show horizontal alignment of support cells in primI neuromasts and a loss of alignment in primII-derived neuromasts as characteristic for single *MZwnt11f1* mutants (Fig. 5k-k’; l-l’). These results indicate that, while PCP is not required for support cells to acquire coordinated organization, the Wnt pathway genes are essential for the proper coordinated organization of support cells in primII-derived neuromasts.

### Wnt pathway genes likely act in the pre-migratory primordium to establish the coordinated support cell organization observed at 5dpf

Because *wnt11f1* is not expressed in 5dpf primII-neuromasts, which show a phenotype in *MZwnt11f1* mutants (Fig. 1 and Fig. 5), we hypothesized that *wnt11f1* might be required earlier in lateral line development to establish hair cell orientation. We therefore tested if Wnt pathway members are either expressed in primII or in adjacent tissues during migration or placode specification stages. At 50 hours post fertilization (hpf), while primII is migrating and has not yet deposited its first neuromast, *vangl2* is expressed in primII, and in primI and primII-derived neuromasts, suggesting that it is acting within lateral line cells (Fig. 6a). As *scrib* is also a core PCP gene it should show the same expression pattern as *vangl2*. However, *scrib* RNA is not detected in the lateral line by in situ hybridization, even though it is expressed at a low level in a previous RNA-Seq analysis ((Jiang et al., 2014), Fig. 6b). In contrast to *vangl2, wnt11f1* mRNA is not detectable in either the migrating primII, or primI-derived neuromasts (Fig. 6c). However, it is expressed in the underlying muscle along the myoseptum (Gordon et al., 2012; Jing et al., 2009), suggesting that the muscle could signal to the neuromasts (Fig. 6d; Fig. Suppl. 10a). In contrast to the *wnt11f1* ligand, the Wnt co-receptor *gpc4* is expressed in the migrating primII and primI- and primII-derived neuromasts, even though its loss only affects primII-derived neuromasts (Fig. 6e, Fig.1). The expression of the *fzd7a* and *fzd7b* receptors is more complex. Both are expressed in the lateral line, but while *fzd7a* is expressed in primII, and primII-derived neuromasts (Fig. 6f), *fzd7b* is also expressed in primII but only primI-derived neuromasts (Fig. 6g). As only *fzd7a* is expressed in the migrating primII but only *fzd7a/7b* double mutants affect primII-derived neuromasts, we hypothesize that Wnt signaling is already acting during pre-migratory stages.

**Figure 6.**
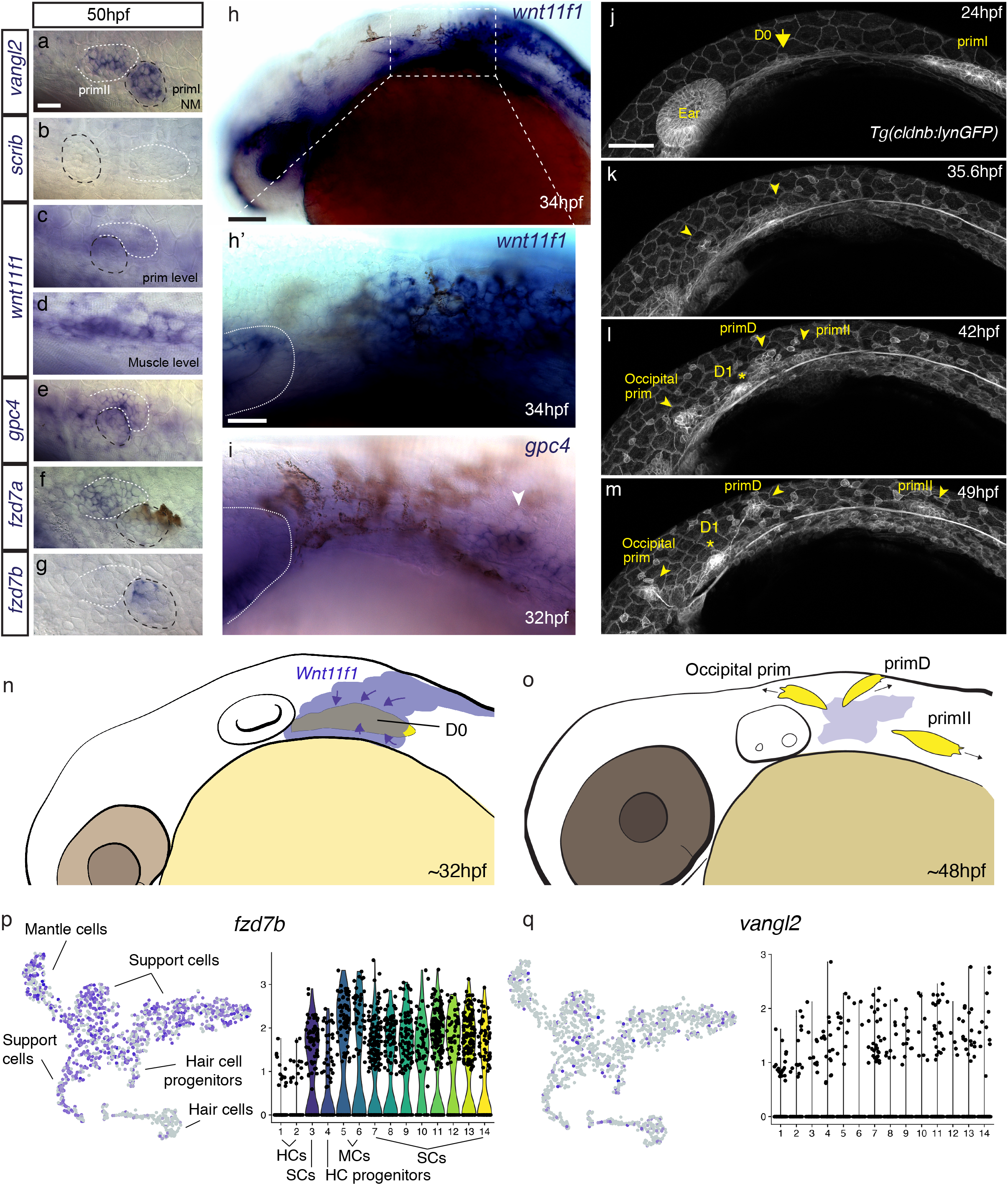
Temporal dynamic expression of the PCP and Wnt pathway genes during formation and migration of primII. a-g. *In situ* localization on 50hpf wild type fish of mRNA for *vangl2* (b) and *scrib* (b), *wnt11f1* (c, at the level of the primordium; d, at the level of the underlying muscle), *gpc4* (e), *fzd7a* (f), *fzd7b* (g). The first neuromast derived from primI is outlined using black, while primII is outlined in white in a-g. h. *In situ* localization of *wnt11f1* mRNA on the anterior region of a 34hpf wild type embryo. h’. Detail of the expression in the area posterior to the ear in h. The ear is delimited by a dashed line. i. *In situ* localization of *gpc4* mRNA in the area posterior to the ear at 32hpf. The ear is delimited by a dashed line. Arrowhead indicates the putative localization of the D0 placode. j-m. Still frames of the time lapse analysis of the formation of primII, primD, occipital prim and D1 labeled in a *Tg(cldnb:lynGFP)* transgenic wild type fish (see also Suppl. Movie 5). Arrow in j indicates the original group of cells that originates the primordia that show primII-polarity. Arrowheads in k-m indicate the different primordia formed. Asterisk in l-m indicates the position of the D1 neuromast. n-o. Schematic cartoon of the proposed mechanism by which Wnt11f1 would signal to the cells in D0 placode (n) and establish support cell organization on the primordia originated from it (o). p. t-SNE plots and violin plots showing expression of *fzd7b* (p) and *vangl2* (q) in a 5dpf neuromast during homeostasis. HCs= Hair Cells, SCs= Support Cells, MCs= Mantle Cells. Scale bar in a= 10μm, h=50μm, h’=25μm, j= 50μm.

Analysis of the expression pattern of the Wnt pathway genes at a time point in which primII is being specified (32hpf) shows that *wnt11f1* is expressed in a dispersed group of cells posterior to the ear immediately adjacent to the forming, *gpc4*-expressing D0 placode (Fig. 6h-h’, Fig. 6i arrowhead). In addition, *wnt11f1* is expressed in the lens, brain, ear and dorsal spinal cord. The D0 placode gives rise to primII, primD (that gives rise to the dorsal lateral line) (Ghysen and Dambly-Chaudiere, 2007; Nunez et al., 2009) and the occipital prim (occipital lateral line) (Fig. 6i, arrowhead).

Time-lapse analysis of the formation of primII in wild type fish shows how primII, primD and the occipital prim arise from the D0 placode between 24 and 48hpf (Fig. 6j-m, Suppl. Video 5). Neuromasts in all these three lateral lines possess the same D-V hair cell orientation in wild type fish (Fig. Suppl. 11a, c) and also show concentric hair cell organization in the Wnt pathway mutants (Fig. Suppl. 11e, g). Due to the fact that the neuromasts derived from primD and the occipital prim do not migrate over any *wnt11f1*-expressing muscle cells, unlike primII in the trunk, we conclude that the hair cell defect in primII neuromasts is not due to loss of Wnt signaling from muscle cells along the horizontal myoseptum. The only time these three lateral lines are all exposed to *wnt11f1* is when they arise from D0, suggesting that *wnt11f1* is required in the placode before the three primordia migrate into different directions (Fig. 6n,o).

Single Cell RNASeq analysis of neuromasts also suggests that the Wnt pathway genes act in support cells, rather than hair cells ((Mark E. Lush, 2018); https://piotrowskilab.shinyapps.io/neuromast_homeostasis_scrnaseq_2018/). *gpc4* and *fzd7a/b* are robustly expressed in support cells but downregulated as cells differentiate into hair cells (Fig. 6, Fig. Suppl. 10b). *vangl2* and *scrib* are expressed in both support and hair cells (Fig. 6, Fig. Suppl. 10b), however the loss of *vangl2* only affects the hair cells. The role of *vangl2* in support cells is therefore still unresolved.

Overall, these results suggest that while PCP genes are expressed and required to establish proper hair cell planar polarization within all neuromasts, Wnt pathway genes act earlier in support cells during D0 placode formation, prior to the differentiation of hair cells. Our data shows that Wnt pathway genes are involved in establishing support cell orientations that secondarily influence hair cell orientations in 5dpf primII neuromasts (Fig. 6n,o). Thus, the loss of Wnt signaling early in development has effects on organ formation several days later.

## DISCUSSION

Several studies have reported that the PCP and Wnt pathway genes are involved in the establishment of hair cell orientation in vertebrates (Dabdoub et al., 2003; Deans et al., 2007; Duncan et al., 2017; Jones et al., 2014; Montcouquiol et al., 2003; Montcouquiol et al., 2006; Qian et al., 2007; Sienknecht et al., 2011; Stoller et al., 2018; Wang et al., 2005). However, the Wnt pathway mutants are often classified as PCP signaling mutants due to the fact that both pathways cause disruptions in the stereotypic arrangement of hair cells. Here we uncovered differences in hair cell phenotypes in zebrafish mutants of the Wnt and PCP signaling pathways. This work demonstrates the power of the lateral line as a model to dissect the roles of these pathways during the morphogenesis of a complex mechanosensory organ that possesses planar polarized hair cells and our data raise a number of interesting issues.

### PCP and Wnt signaling pathways control two different aspects of hair cell orientation

Previous reports have described the different levels of planar organization that hair cells display in the inner ear (reviewed in (Axelrod, 2008; Deans, 2013)). For instance, the term ‘Subcellular Planar Polarity’ refers to the asymmetric position of the kinocilium on one side of the apical surface and stereociliary bundle polarity, while ‘PCP’ refers to the coordinated orientation of neighboring cells. The third level, ‘Tissue Polarity’ refers to the subdivision of hair cells into two groups patterned around the Line of Polarity Reversal in the vestibular system. Since in both PCP and Wnt pathway mutants the kinocilium migrates to the cell periphery (Fig. 1), we assume that the Subcellular Planar Polarity is not affected and the hair cell defects are at the PCP or Tissue Polarity level.

Wnt ligands play different roles in different PCP contexts. For example, Wnt ligands instruct PCP (Chu and Sokol, 2016; Gao et al., 2011; Gros et al., 2009; Wu et al., 2013) or act as gradients that control hair cell orientation in the inner ear (Dabdoub et al., 2003; Qian et al., 2007). Our results demonstrate that in the context of neuromasts, the Wnt pathway genes do not affect PCP signaling as Vangl2 localization is normal in Wnt pathway mutants (Fig. 3). In addition, in *MZwnt11f1;vangl2* double homozygous neuromasts, hair cell orientations are randomized as in *vangl2* single mutants, suggesting that PCP signaling is normal in *MZwnt11f1* mutants (Fig. 2). Recent reports show that Emx2 and Notch signaling control stereocolia bundle orientation and that Vangl2 is normally localized in *emx2* mutants suggesting that this pathway acts in parallel to the PCP pathway (Jacobo A, 2018; Jiang et al., 2017). Emx2 is expressed in only one of the pair of sister hair cells and changes its polarity by 180 degrees, thereby causing the pair of hair cells to adopt opposing polarities. As *emx2* affects hair cell polarity without affecting PCP signaling, we wondered if the Wnt pathway genes acted in concert with Emx2 or Notch signaling. However, we did not detect any significant changes in Notch signaling in *MZwnt11f1* mutants (data not shown). Likewise, the number of Emx2-expressing hair cells is normal in *wnt11f1*, even though they are not localized to one half of the neuromasts but are randomly distributed (Suppl. Fig. 12). We therefore conclude that the Wnt pathway genes are not acting in the same pathway with Notch and Emx2 signaling to regulate hair cell orientation. Instead, our data suggest that Wnt pathway genes act in the surrounding support cells (that do not express *emx2*) whose misalignment causes hair cell orientation defects secondarily in the absence of Wnt signaling. Supporting evidence for this interpretation is also provided by the finding that *gpc4* and *fzd7a/b* are strongly expressed in support and not hair cells in neuromasts ((Mark E. Lush, 2018), Fig. 6, Fig. Suppl. 10). Thus, the coordinated alignment of support cells is necessary for hair cells to acquire proper orientation at the Tissue Polarity level.

An unanswered question is why in Wnt pathway mutants the concentric hair cell phenotype arises sequentially as more cells are added. A concentric orientation might represent the most energy-efficient arrangement for hair cells that still possess functional PCP but that are surrounded by misaligned support cells. Our results stress the importance of studying all cell types in an organ when analyzing hair cell phenotypes in vertebrates. Therefore, *gpc4* and *MZfz7a/7b* mutants previously characterized as PCP signaling mutants in other contexts should be re-evaluated as such.

### How are the different orientations in primI and primII-derived neuromasts achieved?

PCP pathway mutations disrupt hair cell orientations in all neuromasts demonstrating that PCP is necessary for coordinated hair cell orientation in sensory organs derived from all primordia. Thus, the mechanisms that control differential A-P and D-V orientations in primI and primII-derived neuromasts are either acting upstream or in parallel to PCP signaling. Lopez-Schier *et al*. proposed that the direction of the migrating primordia or the freshly deposited neuromasts determines the axis of polarity (Lopez-Schier et al., 2004). For example, neuromasts deposited by primII migrate ventrally after deposition (Ledent, 2002; Sapede et al., 2002), which has been correlated with instructing the axis of hair cell polarity with a 90 degree angle with respect to primI-derived neuromasts. However, the adjacent primI-derived interneuromast cells also undergo a ventral migration and form neuromasts with primI-polarity (Ghysen and Dambly-Chaudiere, 2007). Likewise, the D1 neuromast forms from cells left behind by primI and possesses primI polarity (Fig. Suppl. 11b), even though it undergoes a dorsal migration after being formed (Ghysen and Dambly-Chaudiere, 2007). Therefore, the direction of migration is not an indicator of hair cell polarity. In contrast, our data suggest that support cells in different primordia possess a polarity that is set up as they are formed and which instructs hair cell polarity in later forming neuromasts.

Wnt pathway genes are only required for support cell organization in primII-and not primI-derived neuromasts (Fig. 5) raising the question of how primI-derived neuromasts are polarized at the Tissue Level. Support cells in primI neuromasts could be organized by a mechanism that involves tissue tension as described in the skin of mouse (Aw et al., 2016), ciliated epithelia of Xenopus (Chien et al., 2015; Chien et al., 2018) and in Drosophila (Aigouy et al., 2010; Olguin et al., 2011). Alternatively, primI-derived neuromasts may rely on a different subset of Wnt ligands, HSPGs or Fzd receptors to organize their support cells. Indeed, several other Wnt ligands (Jiang et al., 2014; Romero-Carvajal et al., 2015), HSPGs (Venero Galanternik et al., 2015) and Frizzled receptors (Jiang et al., 2014; Nikaido et al., 2013; Sisson and Topczewski, 2009) are expressed in the lateral line or the surrounding tissues that could be acting on primI-derived neuromasts. To date, the analysis of hair cell orientation in zebrafish mutations in two PCP-related non-canonical Wnt ligands, *wnt5b* (*pipetail*) and *wnt11f2* (formerly known as *wnt11/silberblick*, Postlethwait et al., in preparation), revealed no hair cell orientation defects in any of the primordia (Fig. Suppl. 2, and data not shown). Other factors that might be differentially involved in hair cell orientation are HSPGs such as Syndecan4, which interacts with the PCP pathway during the establishment of hair cell polarity in the inner ear of mice (Escobedo et al., 2013) and Frizzled receptors Fzd6 and Fzd3, whose loss causes hair cell orientation defects in mice (Wang et al., 2006). Analysis of hair cell orientation in mutants for these and other Wnt ligands, HSPGs and Fzd receptors would be much informative as to whether primI-derived neuromasts use a different molecular code to orient their hair cells.

### Neuromast polarity is established early but has long-lasting effects on hair cell polarities

Our data suggest that tissue organization in the primordium is set up early during development and influences hair cell orientation in later stages. This mechanism is conceptually analogous to the ratchet effect proposed by Gurdon et al. (Gurdon et al., 1995) and implies that cells have a ‘memory’ of the initial conditions or signals to which they have been exposed that modulates a later response. A similar process has been proposed to be at work during zebrafish gastrulation, where prolonged cadherin-dependent adhesion initiates Nodal signaling and feeds back to positively increase adhesion and determines mesoderm vs endoderm fate (Barone et al., 2017). Additionally, reports from cells ‘remembering’ the spatial geometry or PCP that precedes cell division using the extracellular matrix or tricellular junctions support this hypothesis (Bosveld et al., 2016; Devenport et al., 2011; Thery et al., 2005). This ‘memory’ of early polarity establishment might explain why primII-derived Wnt pathway mutant neuromasts possess a phenotype, even though *wnt11f1* is not expressed in surrounding tissues at that stage (Fig.1, Fig. 5).

### A PCP-independent mechanism for support cell organization

Our data demonstrate that Wnt pathway genes are required for coordinated alignment of support cells. Rather than acting as an instructive cue, Wnt pathway genes might be acting as permissive factors by controlling cell-cell adhesion. For example, *wnt11f1* destabilizes apical cellcell junctions in the epithelium during pharyngeal pouch formation in zebrafish (Choe et al., 2013) and expression of dominant negative Wnt11 in Xenopus shows that Wnt11 controls cell adhesion through cadherins (Dzamba et al., 2009). Interestingly, *wnt11f2* interacts with *gpc4* and *fzd7a/7b* and influences E-cadherin dependent cell-cell adhesion and cell contact persistence, rather than PCP during zebrafish convergent extension and eye formation (Cavodeassi et al., 2005; Petridou et al., 2018; Ulrich et al., 2005). However, even though *wnt11f1* binds to the same receptors as *wnt11f2, wnt1112* does not affect hair cell development, suggesting that the the two *wnt11* orthologs likely control different adhesion molecules in the different tissues. This interpretation is supported by our finding that transplantated *cdh1* and *cdh2* mutant or morphant neuromast cell into wild type embryos do not possess hair cell orientation defects (data not shown). In search for other adhesion molecules, we also tested a mutation in EpCAM, a major epithelial adhesion molecule expressed in the lateral line and whose function is critical during epiboly (Slanchev et al., 2009). Its loss also does not produce misorientation of the hair cells (data not shown).

Another intriguing possibility is that the Wnt pathway genes might act through the Fat-Dachsous (Ft-Ds) pathway, a possibility not discussed in the literature thus far. In flies, disruptions in Ft-Ds signaling cause swirling wing hair patterns and disruptions of the global alignment of PCP proteins without affecting their asymmetric distribution, reminiscent of the phenotypes we observed in Wnt pathway mutants (Casal et al., 2018; Casal et al., 2006; Devenport, 2014; Gubb and Garcia-Bellido, 1982; Matis and Axelrod, 2013; Thomas and Strutt, 2012). Furthermore, a recent report shows that the Ft-Ds pathway directs the uniform axial orientation of cells in the Drosophila abdomen (Mangione and Martin-Blanco, 2018), which may be similar to the coordinated organization observed in support cells in our study. Interestingly, mutations in Ft-Ds family members disrupt hair cell and cochlear convergent extension in the mouse (Mao et al., 2011; Saburi et al., 2012). Analyses of zebrafish *fat* and *dachsous* mutants would be informative, however the presence of four *fat* genes in the lateral line, combined with the possibility that they act redundantly makes their functional analysis challenging (Jiang et al., 2014; Steiner et al., 2014). Based on the number of studies cited above it is very likely that *wnt11f1, gpc4* and *fzd7a/7b* establish support cell organization by controlling the adhesive properties of these cells. However, the presence of a large number of adhesion molecules, and potential for functional compensation, makes functional analyses challenging.

## Conclusions

In summary, we identified and characterized the Wnt pathway genes *wnt11f1, gpc4* and *fzd7a/7b*, and the PCP gene *scrib* as distinct inputs controlling hair cell orientation in the lateral line of zebrafish. The core PCP components *vangl2* and *scrib* control individual hair cell polarity, while the Wnt pathway genes *wnt11f1/gpc4/fzd7a/7b* affect the alignment of support cells, which are the progenitors of the later forming hair cells (Fig. 7). This interpretation is supported by our findings that PCP mutants lack proper hair cell orientation, even though the hair cells are integrated between support cells that show coordinated alignment. On the other hand, in Wnt pathway mutants support and hair cells are misaligned in the presence of functional PCP signaling. Importantly, scRNASeq analysis of neuromasts shows that *gpc4, fzd7a* and *fzd7b* are robustly expressed in support but not hair cells, whereas *vangl2* is also strongly expressed in hair cells. In addition, because the affected support cells in Wnt pathway mutants are normally only exposed to the *wnt11f1* ligand while the primordia are still in the head region, we propose that Wnt pathway genes act early during development before the hair cells appear.

**Figure 7.**
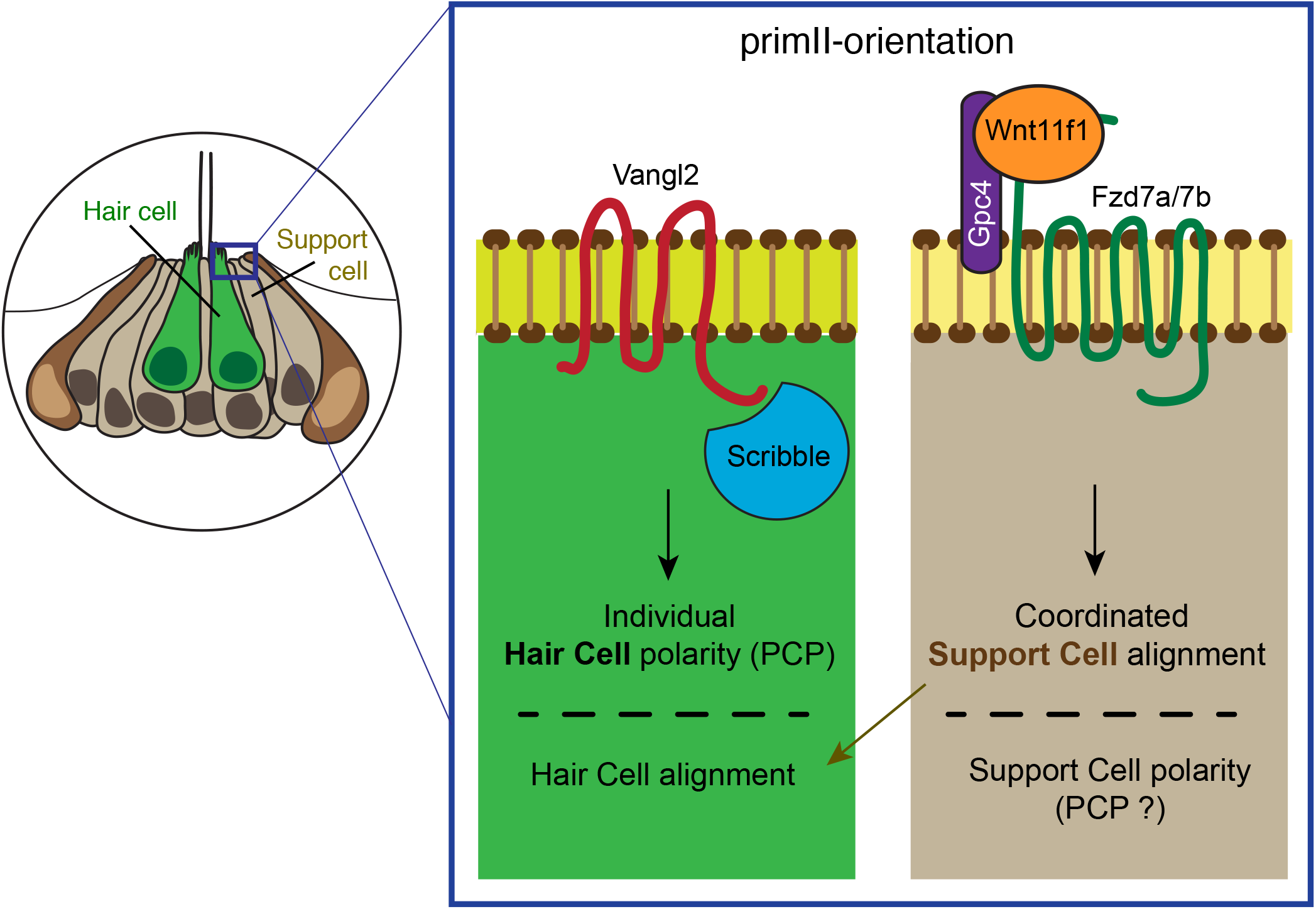
PCP signaling and Wnt pathway act in parallel to establish coordinated hair cell polarity and orientation in the neuromast. The PCP signaling genes Vangl2 and Scrib are required to establish individual polarity in the hair cells (left part of the box), while the Wnt pathway members Wnt11f1, Gpc4 and Fzd7a/7b interact to establish coordinated support cell alignment (right part of the box) in primII-derived neuromasts. The coordinated alignment of support cells, which we assume have normal polarity because coordinated alignment is not disrupted in PCP signaling mutants, will then dictate the individual hair cell orientation.

Our study provides an alternative mechanistic model in which hair cell orientation defects are not only caused by disruptions of the PCP pathway but independently also by other pathways, such as Wnt signaling, which possibly affects adhesion between support and hair cells required to establish or maintain hair cell alignment.

**Figure Supplementary 1 to Figure 1. *wnt11f1* expression in the trunk of a 5dpf larva PrimI-derived neuromasts are labeled using black arrowheads.** primII-derived neuromasts are labeled using white arrowheads and outlined using a dotted line. Note how primI-derived neuromasts show *wnt11f1* mRNA expression on the anterior side and the signal is absent in primII-derived neuromasts (see also Fig. 1d). Scale bar=50μm

**Figure Supplementary 2 to Figure 1. Quantification of the alignment of hair cells with respect to the nearest neighbor in the different conditions analyzed.** a. Diagram showing how the angle between two neighboring hair cells was calculated. Cell 1 is closer to Cell 2, and Cell 2 is closer to Cell 3. The angle with respect to their axes is calculated for each pair. For more details, see Materials and Methods. b-q. Histograms of angular data and fitted distribution (red) for each of the conditions analyzed. b-i. Binned angle distribution in primI-derived neuromasts. b. Distribution in primI-derived neuromasts of wild type fish. Note the tigh degree of alignment between neighbors (Uniform distribution p-value=7.75e-107) and the good fit of the von Mises distribution (R^2^=0.999). c. Distribution in primI-derived neuromasts of Wnt pathway mutants. They all show a high degree of alignment (Uniform distribution p-value in c= 1.21e-277, d= 1.97e-184 and e=5.46e-44) and a good fit for the von Mises distribution (R^2^= 0.999, 0.996 and 0.991, respectively) f. Angle distribution in primI-derived neuromasts of *vangl2* mutants. Neighbor hair cells do not show angle coordination (Uniform distribution p-value= 0.212). g. Angle distribution in primI-derived neuromasts of *MZscrib* mutants. Hair cells show a high degree of coordination between neighbors (Uniform distribution p-value= 7.11e-86), but the distribution is more dispersed than the one shown by wild type fish. h-m. Binned angle distribution in primII-derived neuromasts. h. Angle distribution in primII-derived neuromasts of wild type fish. Note the high degree of alignment between neighbor hair cells (Uniform distribution p-value= 2.08e-106). i-k. Angle distribution in primII-derived neuromasts of Wnt pathway mutants. In all three conditions, hair cells show significant degree of coordination between neighbors (Uniform distribution p-value in i= 3.09e-18, j= 1.97e-07, k= 7.03e-14) and a good fit in the von Mises distribution (R^2^ in i= 0.971, j= 0.892, k= 0.725). l. Angle distribution in primII-derived neuromasts of *vangl2* mutants. Neighbor hair cells do not show coordination in their alignment (Uniform distribution p-value= 0.059). m. Angle distribution in primII-derived neuromasts of *MZscrib* mutants. Hair cells show a high degree of coordination between neighbors (Uniform distribution p-value= 1.09e-33), but the distribution is more dispersed than the one shown by wild type fish. n. Hair cell orientation is disrupted only in primII-derived neuromasts of 5dpf *MZwnt11f1* homozygous siblings and *MZwnt11f1;gpc4* double homozygous fish. o. Hair cell orientation in 5dpf *wnt11f2* homozygous, *wnt11f1* homozygous and double *wnt11f2;wnt11f1* homozygous fish. Note how only a mutation in *wnt11f1* disrupts hair cell orientation in primII-derived neuromasts. The cell axis defined by the position of the kinocilum is labeled in yellow. Scale bar in n and o=5μm.

**Figure Supplementary 3 to Figure 2. Quantification of the alignment of hair cells respect to the nearest tangent (concentricity) in the Wnt pathway mutants *gpc4* and *MZfzd7a/7b* and the PCP mutants *MZscrib*.** a. Diagram showing how the concentricity of the hair cells in each of the conditions was calculated. The angle between a cell’s axis (C) with respect to the tangent (T) of the nearest fitted ellipse is calculated for each cell. For more details, see Materials and Methods. b-d. Phalloidin images showing the cell polarity axis (yellow lines) in primI-derived neuromasts of *gpc4* (b), *MZfzd7a/7b* (c) and *MZscrib* mutants (d). Note that from the distribution of angles with respect to the nearest ellipse tangent (concentricity) none of the mutants shows significant concentricity (Uniform distribution p-values in b= 0.95, c= 0.666, d=0.165; *gpc4* n=222 hair cells, *MZfzd7a/7b* n=74, MZscrib n=535). e-g. Phalloidin images showing the cell polarity axis (yellow lines) in primII-derived neuromasts of *gpc4* (e), *MZfzd7a/7b* (f) and *MZscrib* (g) mutants. For each of the conditions, the distribution of angles with respect to the nearest ellipse tangent (concentricity) for primII-derived neuromasts is shown. Only the Wnt pathway mutants show significative concentricity (Uniform distribution p-values in e= 2.03e-11, f= 2.05e-12, g=0.488; *gpc4* n=214 hair cells, *MZfzd7a/7b* n=182, *MZscrib* n=392). Additionally, the Von Mises distribution p-value and the R^2^ fit were calculated for each of the conditions, and both Wnt pathway mutants show a good fit in primII-derived neuromasts (R^2^ for e= 0.98, f= 0.971). Yellow lines in b-g indicate the hair cell polarity axis, determined by the position of the kinocilium. Not Concentric vs Concentric labels were based on calculations in b-g. Scale bar in b= 5μm.

**Figure Supplementary 4 to Figure 2. Quantification of the alignment of hair cells with respect to the nearest neighbor in *MZwnt11f1* and *MZwnt11f1;vangl2* mutants.** a-b. Angle distribution in primI-derived neuromasts of single *MZwnt11f1* siblings (a) and *MZwnt11f1;vangl2* mutants (b). Note how the high degree of alignment in a (Uniform distribution p-value= 1.82e-223) is lost in b (Uniform distribution p-value= 0.0395). c-d. Angle distribution in primII-derived neuromasts of single *MZwnt11f1* siblings (c) and double *MZwnt11f1;vangl2* mutants (d). Note how the high degree of alignment in c (Uniform distribution p-value= 1.29e-14) is lost in d (Uniform distribution p-value= 0.069).

**Figure Supplementary 5 to Figure 3. Vangl2 enrichment respect to the position of the kinocilium** From all the hair cells that show asymmetric Vangl2, we measured the pole of enrichment with respect to the position of the kinocilium. The results are categorized into three possibilities: asymmetric on the same pole as the kinocilium, asymmetric on the opposite pole than the kinocilium, and asymmetric but outside of the axis defined by the kinocilium. We observe differences between *MZscrib* and both wild type and *MZscrib* siblings (Fisher’s exact test p-val WT primI vs *MZscrib* primI= 0.09724; p-val *MZscrib* siblings primI vs *MZscrib* primI= 0.08285; p-val WT primII vs *MZscrib* primII= 0.02568; p-val *MZscrib* siblings primII vs *MZscrib* primII= 0.03557). Wild type and *MZwnt11f1* mutants do not show differences (p-val WT primI vs *MZwnt11f1* primI= 1; p-val WT primII vs *MZwnt11f1* primII= 0.439).

**Figure Supplementary 6 to Figure 3. Vangl2 antibody staining is absent in *vangl2* mutants.** Vangl2 antibody staining on *vangl2* siblings primI (a) and primII (b). Vangl2 antibody staining on primI (c) and primII (d) of *vangl2* mutants. Scale bar= 5μm.

**Figure Supplementary 7 to Figure 4. Analysis of cell behaviors during the development of the hair cells in primII-derived neuromasts** a-b. Visual representation of the individual time lapses of hair cell formation in primII-derived neuromasts of wild type (a) and *MZwnt11f1* mutants (b). Each hair cell precursor (yellow) divides into two hair cells (red and blue) and either rearrange or not. c. Duration of the rearrangement that the hair cell progenitors undergo after division in wild type and *MZwnt11f1* mutant embryos. d. Percentage of hair cell progenitors that undergo rearrangements after division in wild type and *MZwnt11f1* mutant fish. e. Division angle with respect to the radius of each division event in hair cell progenitors of wild type and *MZwnt11f1* mutant fish, and the visual representation of each division event (respect to the XY axis).

**Figure Supplementary 8 to Figure 4. Quantification of the alignment of hair cells during the development of primII in *vangl2* and *MZwnt11f1* mutant fish at 4 and 5dpf.** Histograms of angular data and fitted distribution (red) for each of the conditions analyzed. Note how none of the conditions except for *MZwnt11f1* at 5dpf show coordination on the alignment of their hair cells with respect to their neighbor (Uniform distribution p-values for *vangl2* at 4dpf=0.733, *vangl2* at 5dpf=0.512, *MZwnt11f1* at 4dpf=0.066, *MZwnt11f1* at 5dpf=1.46e-04).

**Figure Supplementary 9 for Figure 5. Pipeline for the segmentation of Phalloidin stainings of neuromasts 3 hours after hair cell ablation.** Each Phalloidin image is sharpened and segmented. After segmentation, only the support cells are kept and used for fitting the Ellipse that determines the cell’s long axis. For further information, please see Materials and Methods.

**Figure Supplementary 10 to Figure 6. *wnt11f1 in situ* on a 50hpf wild type fish and single-cell analysis of RNA expression of *fzd7a, gpc4* and *scrib*.** a. Low magnification picture of *wnt11f1* mRNA expression on the trunk of a 50hpf fish. *wnt11f1* is expressed along the myoseptum. b. t-SNE plots and violin plots showing expression of *fzd7a, gpc4* and *scrib* in a 5dpf homeostasis neuromast. Scale bar in a= 50μm.

**Figure Supplementary 11 to Figure 6. Neuromasts in the Occipital and Dorsal lateral lines show hair cell orientation defects in *MZwnt11f1* mutant fish.** a-c. Hair cell orientation of neuromasts in wild type. e-g. Hair cell orientation of neuromasts in *MZwnt11f1* mutants. Neuromasts that show a hair cell phenotype in *MZwnt11f1* mutants are labeled as Concentric. D1=Dorsal 1. D2= Dorsal 2 (belonging to the Dorsal lateral line). Scale bar in a= 5μm.

**Figure Supplementary 12. Distribution, but not ratio of expression, of Emx+ hair cells is affected in primII-derived neuromasts of *MZwnt11f1* mutants.** Antibody staining for the transcription factor Emx2 in 5dpf wild type (a-b) and *MZwnt11f1* mutants (c-d). In wild type, Emx+ hair cells are biased towards the anterior pole of the primI-derived neuromasts (a’, p=val 7.6e-6) and towards the dorsal pole in primII-derived neuromasts (b’, p-val=7e-4). In *MZwnt11f1* mutants, the distribution is still biased in primI-derived neuromasts (c’, p-val=8.5e-10) but it is disrupted in primII-derived neuromasts (d’, p-val=0.2). Additionally, in both wild type and *MZwnt11f1* mutants, approximately half of the hair cells express Emx2 (e). The arrows in a’, b’ and c’ indicate the direction of bias in each case. *** denotes statistical significance for biased location. N.S= not significant.

## Supplementary Video 1 to Figure 4

Time lapse analysis of hair cell formation during primII migration of a wild type *Tg(cxcr4b:H2A-GFP)* fish. The hair cell progenitor (labeled in yellow) will divide and produce two hair cells (labeled in purple and cyan). Note how the purple cell is in the dorsal pole after division and reverses position with the cyan cell.

## Supplementary Video 2 to Figure 4

Time lapse analysis of hair cell orientation of a 3dpf wild type *Tg(myo6b:β-Actin-GFP)* transgenic fish. The top down view shows the orientation of the actin-rich cuticular plates throughout the duration of the time lapse.

## Supplementary Video 3 to Figure 4

Time lapse analysis of hair cell orientation of a 3dpf *MZwnt11f1* mutant *Tg(myo6b:β-Actin-GFP)* transgenic fish. Note how the orientation of the actin-rich cuticular plates changes over time.

## Supplementary Video 4 to Figure 4

Time lapse analysis of hair cell orientation of a 3dpf *vangl2* mutant *Tg(myo6b:β-Actin-GFP)* transgenic fish. Note how the orientation of the actin-rich cuticular plates changes over time.

## Supplementary Video 5 to Figure 6

Time lapse analysis of the formation of primII, primD, Occipital prim and D1 neuromast on a 24hpf wild type *Tg(cldnb:lyn-GFP)* transgenic fish. The Ear is on the left, and primI will migrate posteriorly (right) out of the field of view at the beginning of the video. The D0 placode is labeled using an arrow, and each primordium subsequently derived is labeled using an arrowhead. The D1 neuromast is labeled using an asterisk.

## MATERIALS AND METHODS

### Fish husbandry

All experiments were performed per guidelines established by the Stowers Institute IACUC review board. The following mutant fish strains previously described were used: *tri^m209^* (Stemple et al., 1996), *gpc4^fr6^* (Topczewski et al., 2001), *wnt11f1^fh224^*(*wnt11r^fh224^* in (Banerjee et al., 2011)), *fzd7a^e3^* and *fzd7b^hu3495^* (Quesada-Hernandez et al., 2010), *scrib^rw468^*(*a* kind gift from C Walsh; (Wada et al., 2005)).

The following transgenic fish lines were used: *tg(cldnb:lynGFP)^zf106^*(Haas and Gilmour, 2006), *tg(myo6b:β-Actin-GFP)(Kindt* et al., 2012), tg(cxcr4b:H2A-GFP)(Kozlovskaja-Gumbriene et al., 2017), *Tg(sqET4)* (Parinov et al., 2004), *tg(cldnB:H2A-mCherry)(Romero-Carvajal* et al., 2015).

### *In situ* hybridization

*In situ* hybridization was performed as described in (Kopinke et al., 2006) with modifications as in (Ma et al., 2008). The following probes were used: *wnt11f1* (*wnt11r* in (Jing et al., 2009), *gpc4* (Venero Galanternik et al., 2015), *fzd7a* and *fzd7b* (Nikaido et al., 2013). The *vangl2* probe was generated using the following pair of Fw/Rv primers (TCATCTCAGAAGCGATCCAGTA / CCACATGCATCAGACCTAAAAA), and the *scrib* probe was generated using the following pair of Fw/Rv primers (AGGATCTTGCCAAGCAAGTG / CCCTGGCGTAGTTTACGTTT) from wild type AB gDNA.

### Immunohistochemistry and Phalloidin staining

For β-II-Spectrin and Vangl2 double antibody staining, embryos were fixed in 2% TCA in water at room temperature for at least 4h. Embryos were then washed briefly in 0.5% Triton-X in PBS (PBSTx) and then blocked for 2h in PBSTx + 2% Normal Goat Serum. Primary antibodies Rabbit anti-Vangl2 (1:100, Anaspec, AS-55659s, now discontinued) and Mouse anti β-II-Spectrin (1:200, BD Transduction, 612562) were diluted in Blocking solution and incubated overnight at 4C. Primary antibody was washed using PBSTx and then detected using an Alexa-568 Anti-Rabbit and Alexa-488 Anti-Mouse Secondary antibodies diluted 1:500 in Blocking Solution, incubated overnight at 4C. Secondary antibody was washed thoroughly using PBSTx before imaging.

For Phalloidin staining, embryos were fixed in 4% PFA in 1X PBS at room temperature for at least 2 hours. PFA-fixed embryos were briefly washed in 0.5% Triton-X in 1X PBS and then permeabilized in 2% PBSTx for 2 hours at room temperature. Subsequently, the embryos were stained with either Alexa-488 or Alexa-568 Phalloidin (1:40 in PBSTx 0.5%) for 2 hours. After staining, the embryos were washed thoroughly using PBSTx 0.5% and imaged.

### Neomycin treatment

To kill hair cells, 5dpf embryos were treated with 300uM Neomycin in 0.5 E2 media for 30 minutes (Harris et al., 2003). Neomycin was then washed away using fresh E2 media, and waited 3h before fixation in 4% PFA in 1X PBS.

### Time-lapse imaging and imaging

Time lapse analysis and imaging of live larvae were performed as described in (Venero Galanternik et al., 2016).

### Hair cell orientation analysis

To calculate the hair cell orientation from the Phalloidin stained neuromasts, the Line tool from FIJI (Schindelin et al., 2012) was used. A straight line was drawn from the pole opposite to the kinocilium towards the pole were the kinocilium was found. To create the Rose diagrams to represent hair cell orientation, angles measured and then plotted using the rose.diag() function in R ggplot. Statistical analysis was performed for each condition against wild type using Fisher’s exact test. To determine A-P or D-V binomial distribution, the data are binned to four groups, left, right, bottom, and up. The binomial test is used to test whether “right+left” counts are different from 50% of total counts.

### Hair cell alignment respect to the neighbor and respect to the nearest tangent (concentricity) analysis

To measure the alignment of the hair cells, we must choose a structure against which to compare the cell’s angles. Specifically, we are looking at relative differences in the angles and an assumed underlying arrangement, rather than comparing them to some absolute feature, such as the horizontal. The first comparison we consider is the angular difference between nearest neighbors. Note although it is possible to provide a consistent polarization of the cell and, thus, measure the angle between two nearest major axes on a scale of (−180, 180)° we are not currently interested in the polarization, thus, we always take the smallest angle between the two major axes. Hence, the angles will be on a scale of (−90,90)°. Further, the sign convention we will be using is positive if the angle measured from the current cell is anticlockwise, and negative otherwise. The second comparison we consider is cell alignment when compared to a fitted ellipse. Initially, we take the cell centers of the Lines calculated in the ‘*Hair cell orientation analysis*’ section, and use a least squares fitting algorithm based on principal component analysis to create an ellipse of best fit. Each cell is then projected to the closest point on the fitted ellipse’s boundary and their major axis angle is compared to that of the ellipse’s tangent at the nearest point. The smallest angle is extracted using the above sign rule. The ‘Nearest neighbor measurement’ provides us with a measurement of how aligned each cell is with its neighbors, whilst the ‘Nearest tangent measurement’ will provide us with a measurement of how aligned each cell is with the fitted ellipse. Specifically, this comparison will suggest whether there is a circular structure underlying the cell alignment. Having derived the data from the experimental results we plot the information as a histogram. Specifically, the range (−90, 90)° is divided up into 12 bins and we tally how often each angle falls within one of these bins. Dividing by the total number of results then normalizes the histograms to represent a probability. Each angle distribution was first tested against the null hypothesis that there was no preferred direction. Thus, we compare the probability histogram against the uniform distribution and see if there was a significant difference, namely p < 0.01. If there was a significant difference we then compared the solutions to the Von Mises distribution, which is the generalized Normal distribution that is periodic on the angular domain. The fitting of the Von Mises distribution tells us if there is a preferred direction and how variable the alignment is.

### Vangl2 asymmetry quantification

Vangl2 asymmetry quantifications from immunolabeled embryos were performed from raw images. Upon visual inspection, Vangl2 being ‘asymmetric’ vs ‘not being asymmetric’ was assigned and quantified using the Cell Counter function of FIJI. Fisher’s exact test was performed to determine statistical differences between samples. From the hair cells that showed Vangl2 asymmetry, a second analysis was performed and classified into three categories: we quantified whether the enrichment was (1) in the pole where the kinocilium is, (2) on the opposite pole, or (3) enriched but out of the axis determined by the kinocilium. Fisher’s exact test was performed to determine statistical differences between samples. The double immunolabeling images were processed for contrast and sharpness only for visual display in the Figure afterwards using Photoshop CS6.

### Generation of the *Tg(myo6b:GFP-XVangl2)* construct

GFP-XVangl2 was amplified from pDest2-GFP-Vangl2 plasmid (A kind gift from J Wallingford, (Butler and Wallingford, 2018)) and Topo TA cloned. The Topo vector was then digested using KpnI and SacI, and ligated into the KpnI-SacI sites of the Tol2 middle entry vector (Kwan et al., 2007) using Quick ligase (NEB). The middle entry GFP-XVangl2 vector was recombined via a Gateway reaction with the 5’ *myo6b* vector (Kindt et al., 2012). The final myo6b:GFP-XVangl2 plasmid was co-injected into AB or *MZwnt11f1* mutant embryos together with *Tol2* mRNA at a final concentration of 30ng/ul.

### GFP-XVangl2 profile analysis

Images were acquired in a Zeiss 780 microscope using a 40x/1.1 W Korr objective. Raw images of Phalloidin stained animals were used for the analysis of GFP intensity. GFP-XVangl2 polarity was determined in a manner analogous to (Smith et al., 2013). These utilized custom plugins as described above. Average cortical line profiles were generated with a 4 pixel thickness utilizing the “polyline kymograph jru v1” plugin. Then maximum angular position of intensity was determined using the “trajectory statistics jru v2” plugin. Profiles were then aligned so that the maximum intensity occurred at angle 0 with “set multi plot offsets jru v1”. Next, angles were wrapped to values between −179 and +180 degrees with “wrap angle profile jru v1”. Finally, profiles were averaged with “average trajectories jru v1” with error bars indicating the standard error in the mean. For some presentations, averaged profiles were normalized to a maximum intensity of 1 for comparison purposes. Peak widths were determined by fitting profiles to a Gaussian peak function via non-linear least squares fitting (plugin is fit multi gaussian jru v1”) with error bars determined by fitting 100 Monte Carlo simulated random data sets (Bevington and Robinson, 2003). Polarity ratios were determined by averaging three values surrounding the 0, 90, and 180 degree data points and calculating their ratios with errors propagated according to Bevington and Robinson, 2003.

### Hair cell progenitor angle of division analysis

Imaging was performed in a Zeiss 780 using a 40x/1.1 W Korr objective, under conditions described above (*Time-lapse imaging* section). To obtain a quantitative measurement of how hair cell progenitors divide within the neuromast, we acquired time-lapses of migrating primII in wild type and *MZwnt11f1* mutant embryos in a *tg(cxcr4b:H2A-GFP); tg(myo6b:β-Actin-GFP*) double background, scanning every 5 minutes. We calculated the division angle of the dividing progenitor with respect to the radius from the center of division, to the center of the neuromast. We used Imaris to manually mark the location of all cell divisions and the center point of the depositing neuromast for all timepoints. To find the center of the neuromast we create a Surface object using the H2A-GFP channel with very coarse surface detail (surface grain size ~ 20μm). For each cell division, we used Spots to manually mark the center of each daughter hair cell. The Surface and Spots data for every timepoint is exported and aggregated in an Excel spreadsheet. With the exported data in Excel, we calculated two vectors for every dividing cell pair. We calculated the vector between each of the daughter cells, as well as the vector from the center of the neuromast to the center of cell division. The smallest angle between the two vectors is calculated to give the division angle with respect to the radius, which is the range from 0-90 degrees. Division angles for wildtype and mutant were then plotted and t-test analysis analysis performed to calculate statistical differences between wild type and *MZwnt11f1* mutants.

### Support cell orientation analysis

To obtain the support cell orientation, Phalloidin-stained neuromasts were processed in FIJI (Schindelin et al., 2012). A custom ‘gray morphology jru v1’ plugin was used to sharpen the signal from the cell membranes and afterwards segmented using the MorphoJ plugin (Klingenberg, 2011). Once a mask of the cells within the neuromast was obtained, we found the orientation of each cell using FIJI’s “Analyze Particles” function. This function fits each cell to an ellipse in order to provide spatial data, including the center point, major and minor axis length, and the major and minor axis angle in cartesian coordinates. We exported the data from FIJI to Python where we processed and plotted the data. We used Matplotlib to create a histogram of the distribution of cell orientation angles in the range of 0-90 degrees. To visualize and verify the data, we also plotted the major axis vector on top of the cell mask image for each sample.

## Acknowledgements

We would like to thank Dr. Robb Krumlauf for comments on the manuscript. We also thank Dr. Robert Duncan for the initial *wnt11f1* in situ analysis, the Aquatics Facility of the Stowers Institute for Medical Research for their outstanding husbandry work. This work was funded by an NIH award RO1 NS082567 to C.M., an NIH (NIDCD) award 1R01DC015488-01A1 to T.P and by institutional support from the Stowers Institute for Medical Research to T.P.

## Author contributions

J.N.A designed the experiments, collected, analyzed and interpreted the data, and wrote the manuscript; R.L.A, T.W, J.R.U, H.L analyzed the data; C.M provided the *wnt11f1^fh224^* mutants; M.G.V. discovered the hair cell orientation phenotype in *wnt11f1* mutants. T.P designed and interpreted the experiments, and wrote the manuscript.

