## Supplementary Figures for "Parallel control of mechanosensory hair cell orientation by the PCP and Wnt pathways"

### Figure Supplementary 1 for Figure 1

*wnt11f1* is only expressed by primI-derived neuromasts

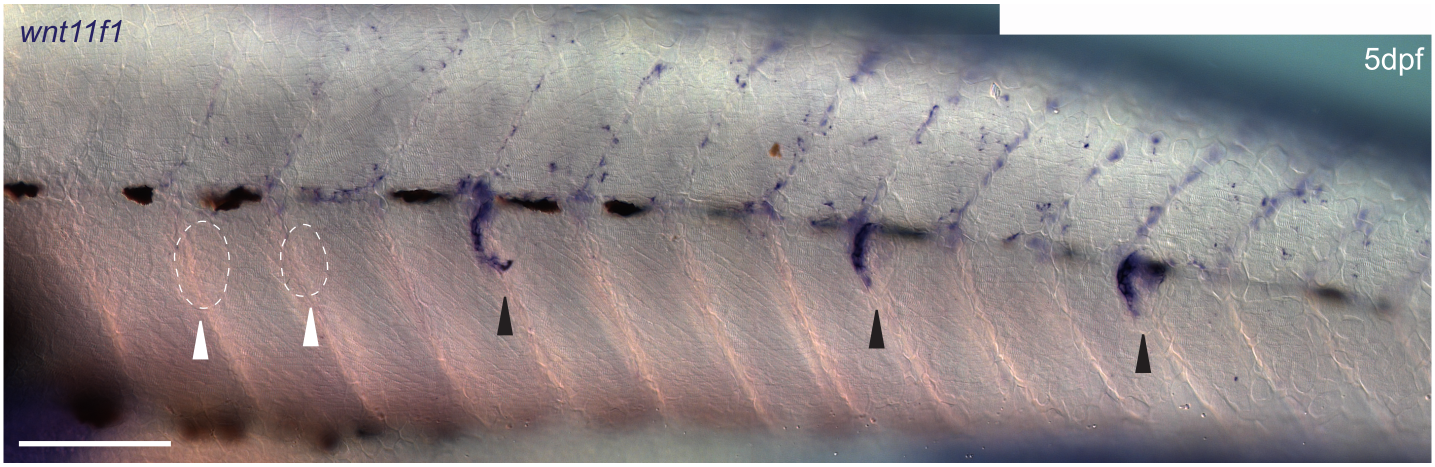

### Figure Supplementary 2 for Figure 1

Analysis of hair cell orientation with respect to the nearest neighbor
*(do hair cells show relative orientation with respect to each other?)*

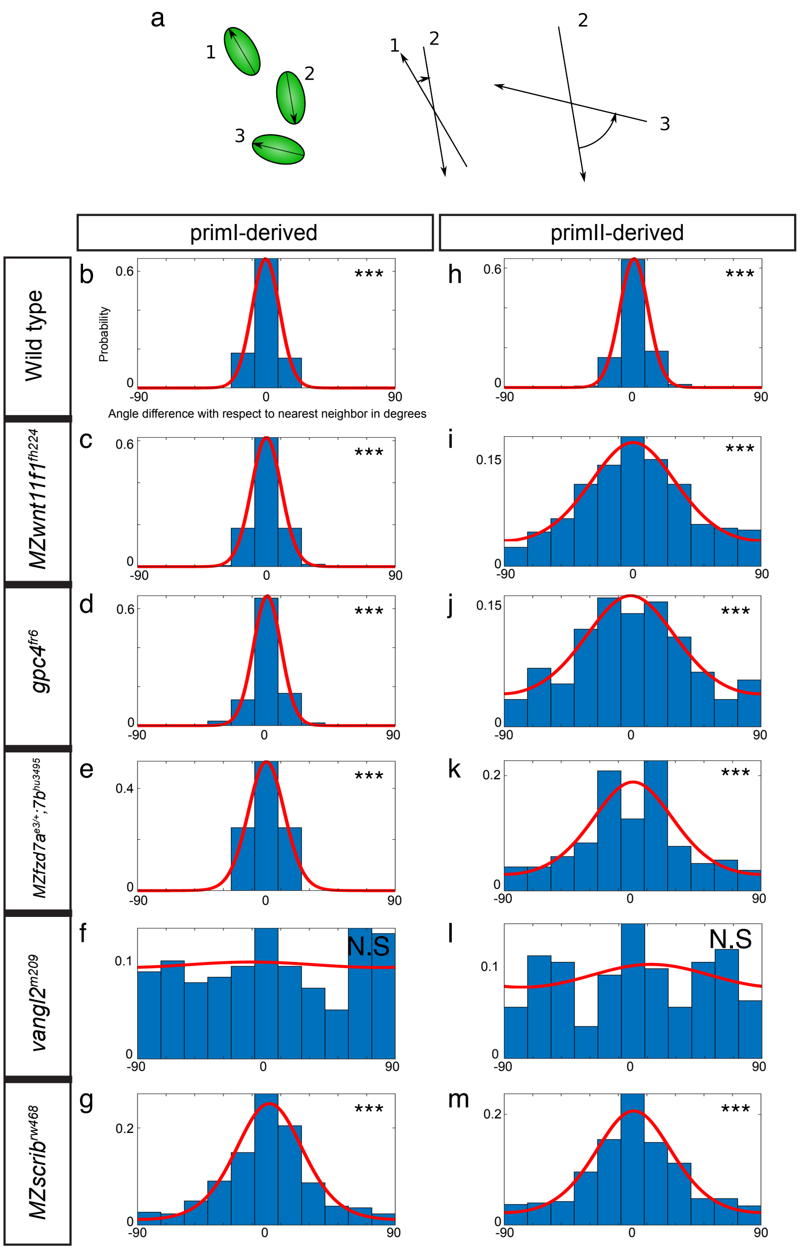


#### Hair cell orientation in double mutants

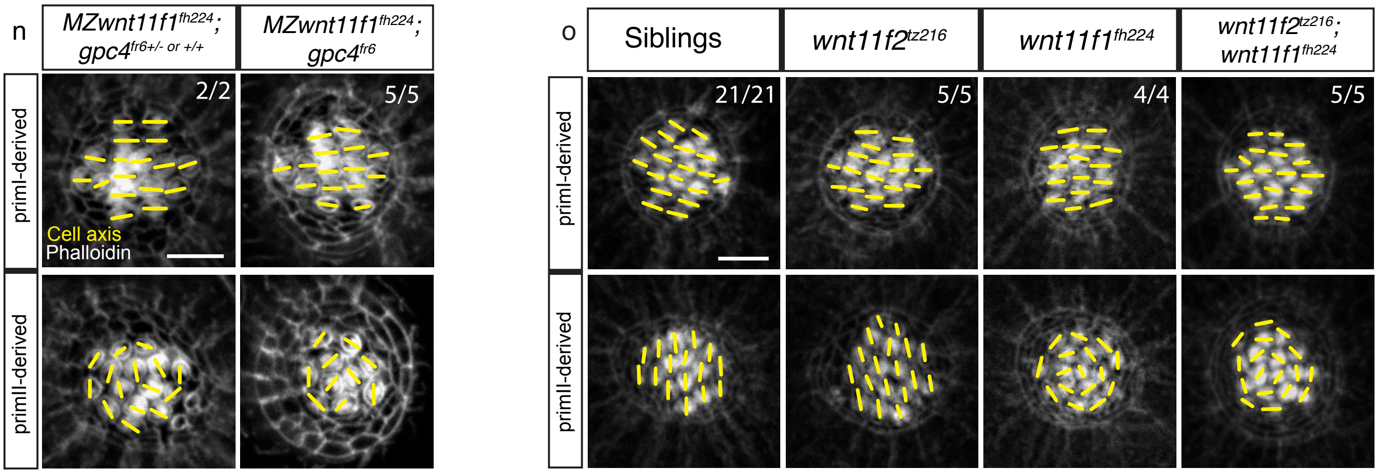

### Figure Supplementary 3 for Figure 2

Analysis of angle difference between a given hair cell's orientation
and the nearest fitting ellipse (concentricity)

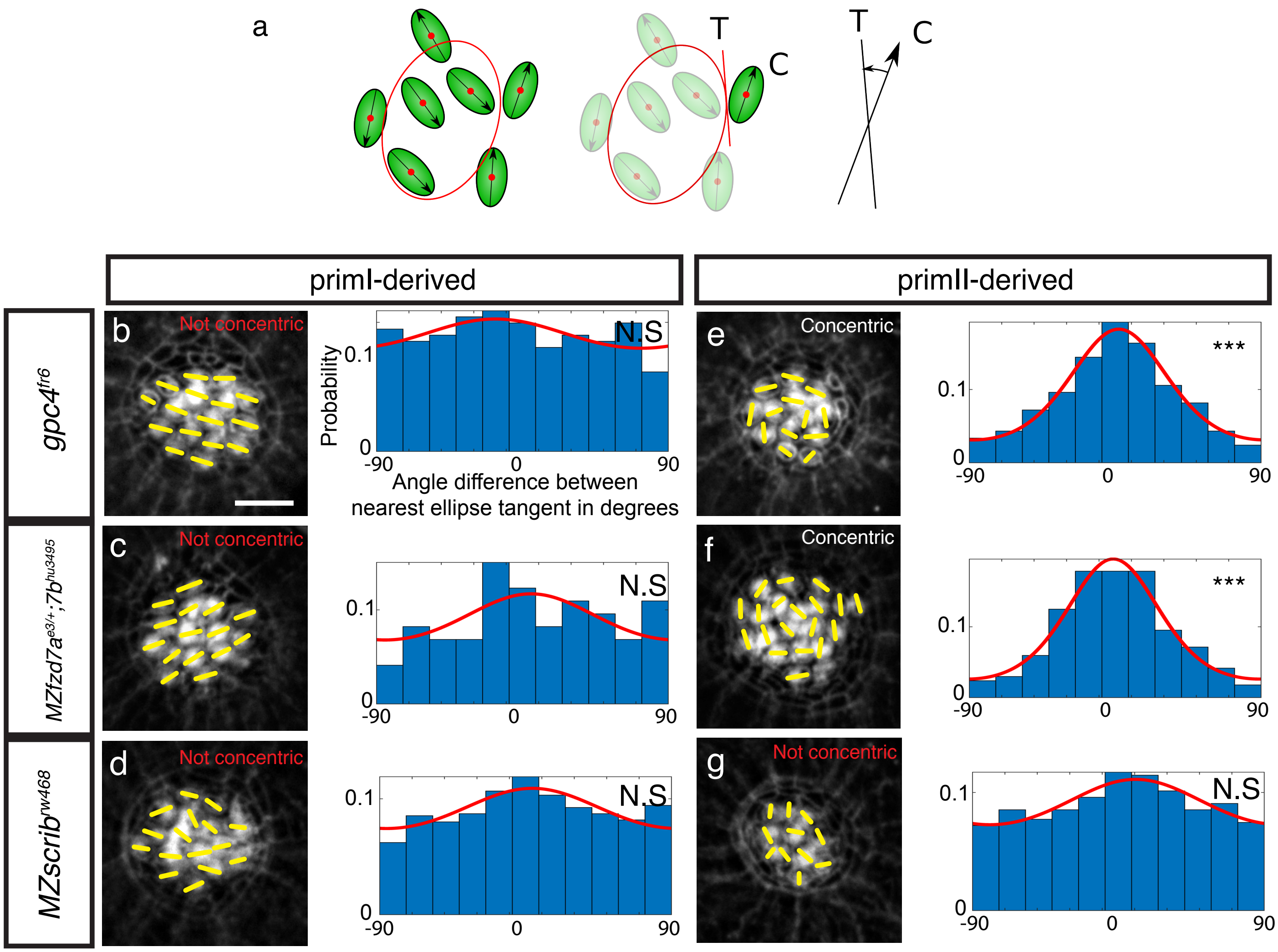

### Figure Supplementary 4 for Figure 2

Analysis of hair cell orientation with respect to the nearest neighbor

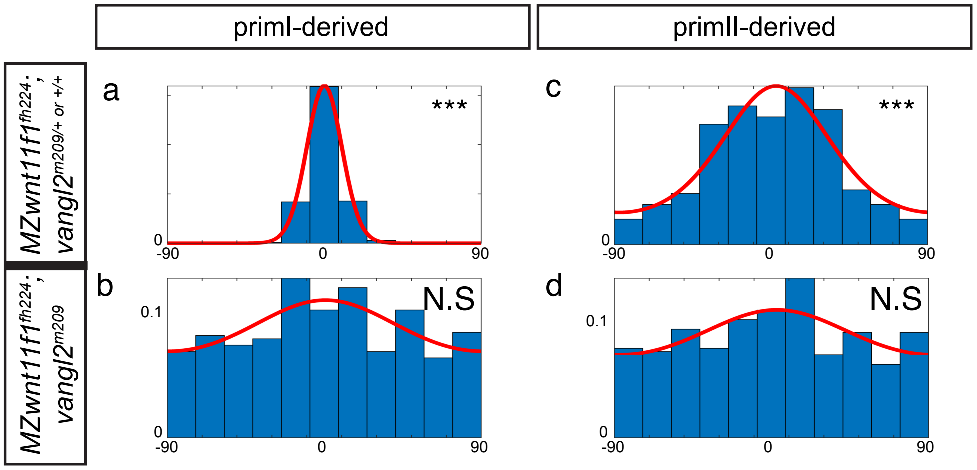

### Figure Supplementary 5 for Figure 3

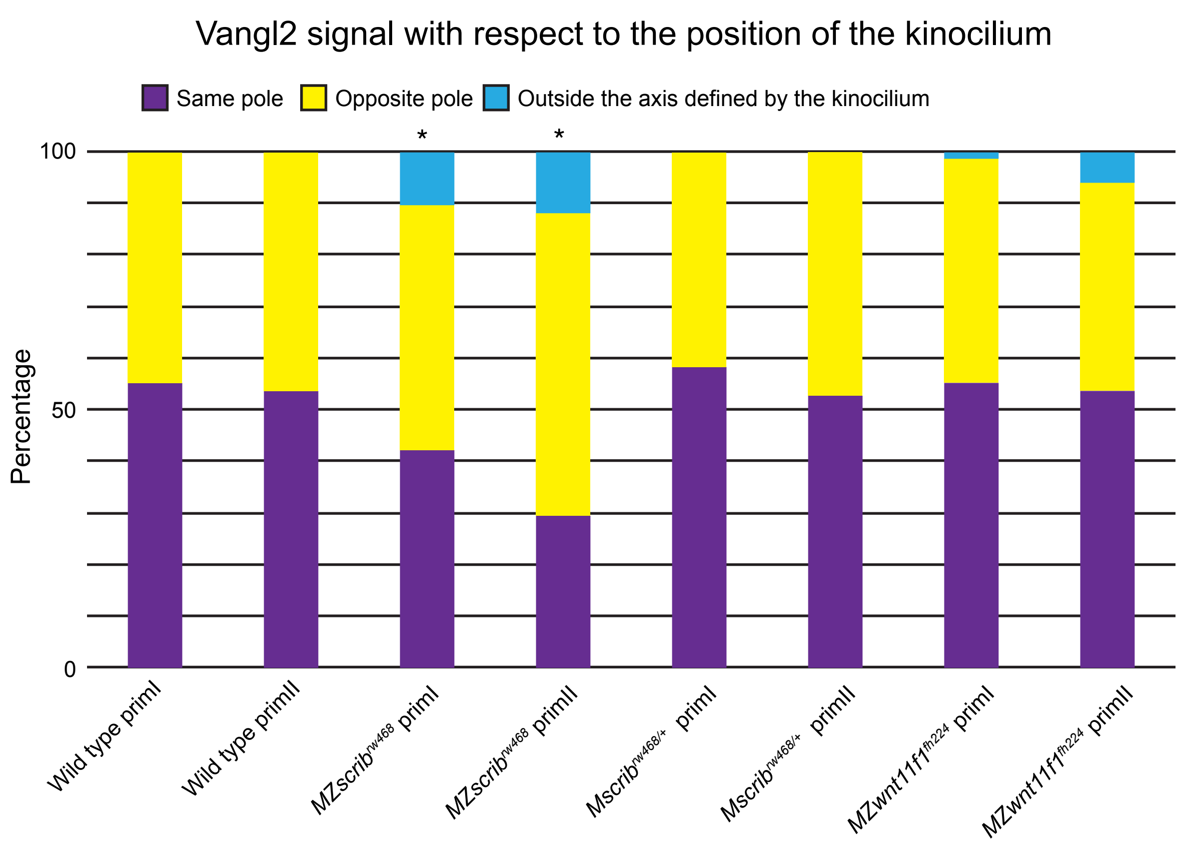

### Figure Supplementary 6 for Figure 3

Vangl2 localization is absent in *vangl2$^{m209}$* mutants

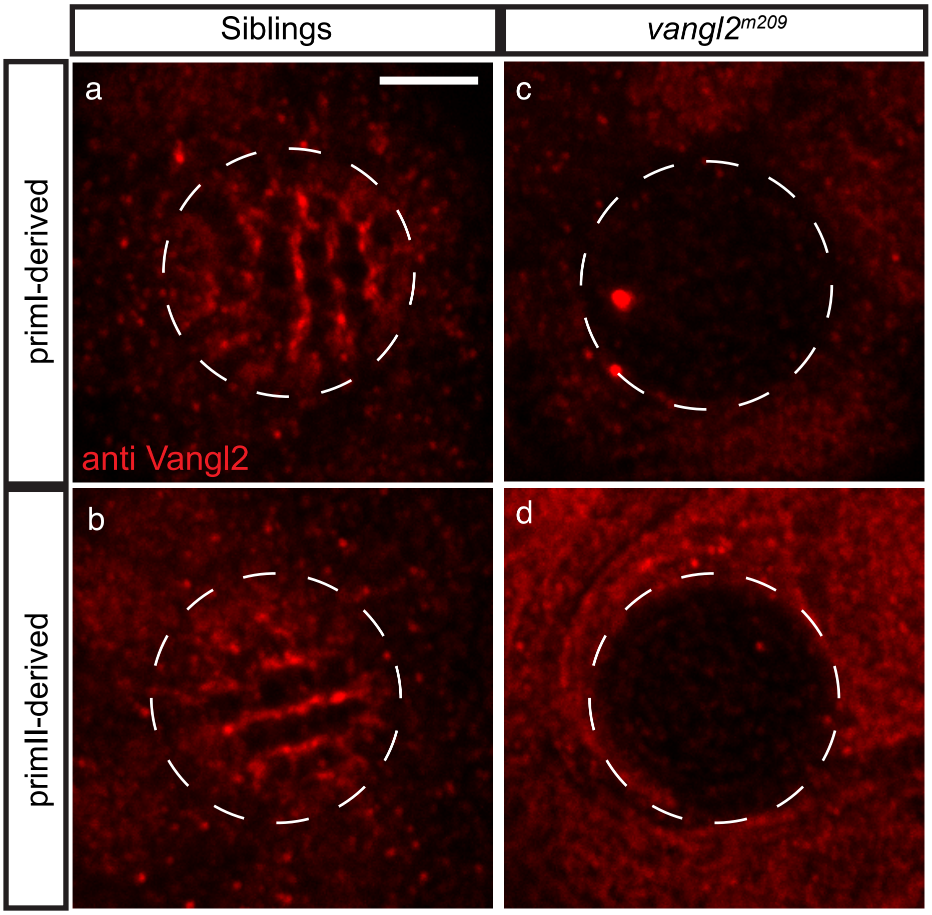

### Figure Supplementary 7 for Figure 4

Analyses of cell behaviors of hair cell precursors during development show no differences between Wild type and *MZwnt11f1* mutants

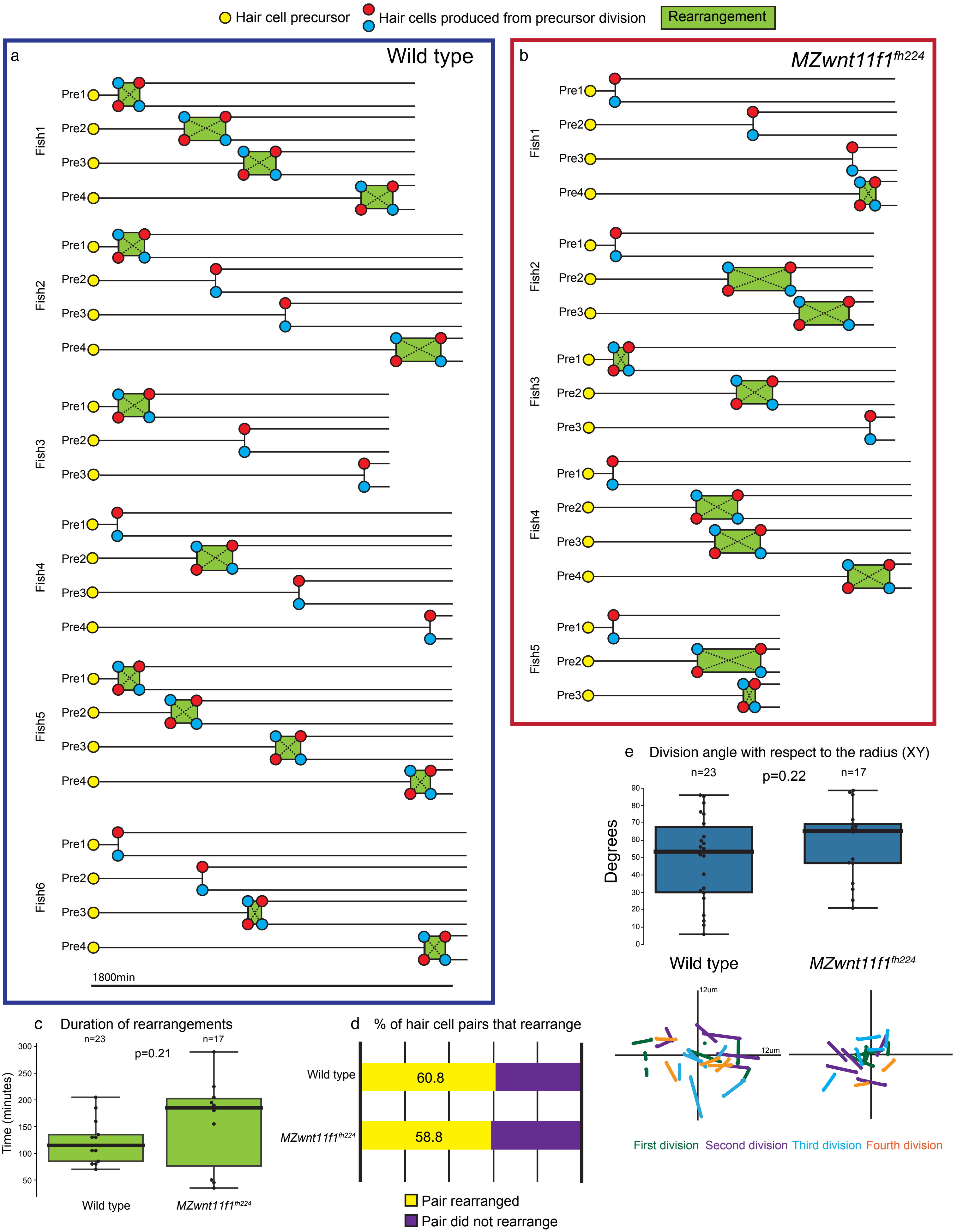

### Figure Supplementary 8 for Figure 4

Angle comparison with nearest neighbor of hair cells in primII-derived neuromasts

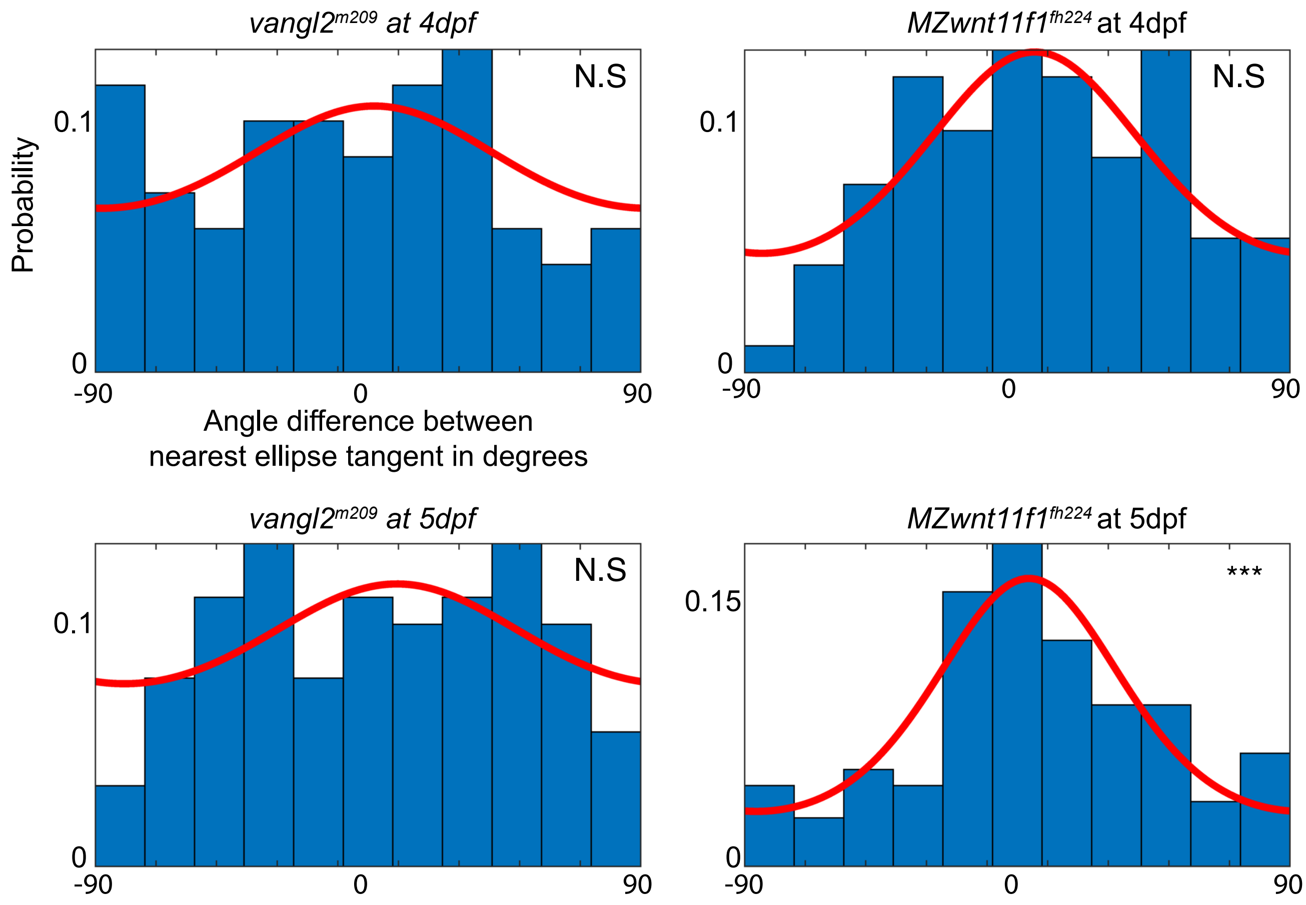

### Figure Supplementary 9 for Figure 5

Pipeline for Segmentation of Phalloidin images for Support Cell orientation analysis

**Step 1**
Phalloidin staining

**Step 2**
Custom sharpening

**Step 3**
MorphoJ segmentation

**Step 4**
Image Clean-up

**Step 5**
Fitting ellipse

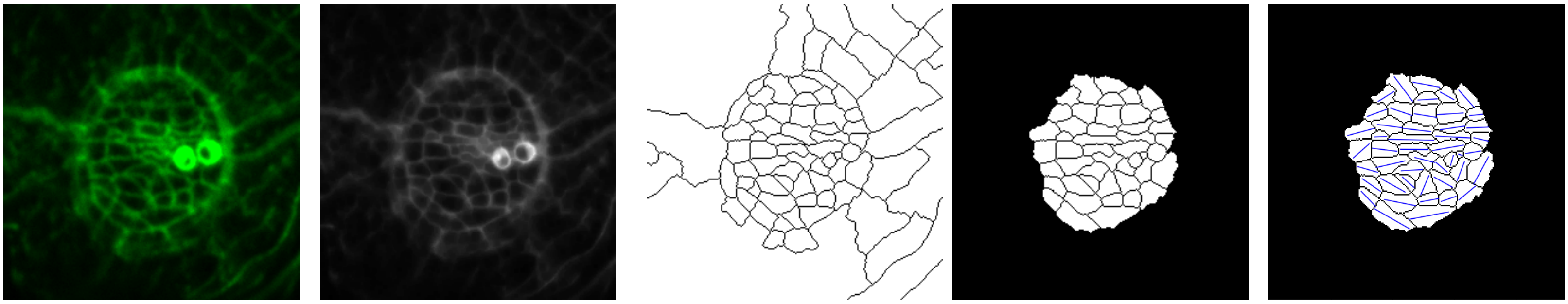

(3h after Neomycin)

### Figure Supplementary 10 for Figure 6

*wnt11f1* is expressed in the muscle

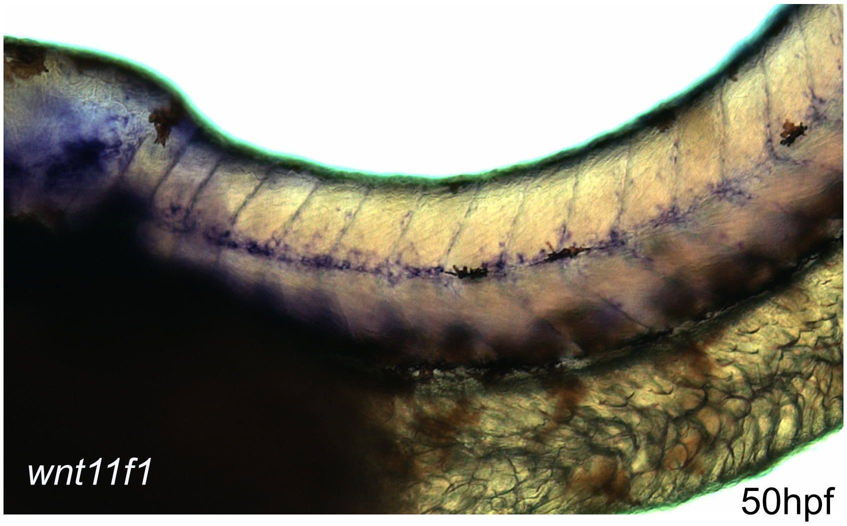


*fzd7a* *gpc4* *scrib*

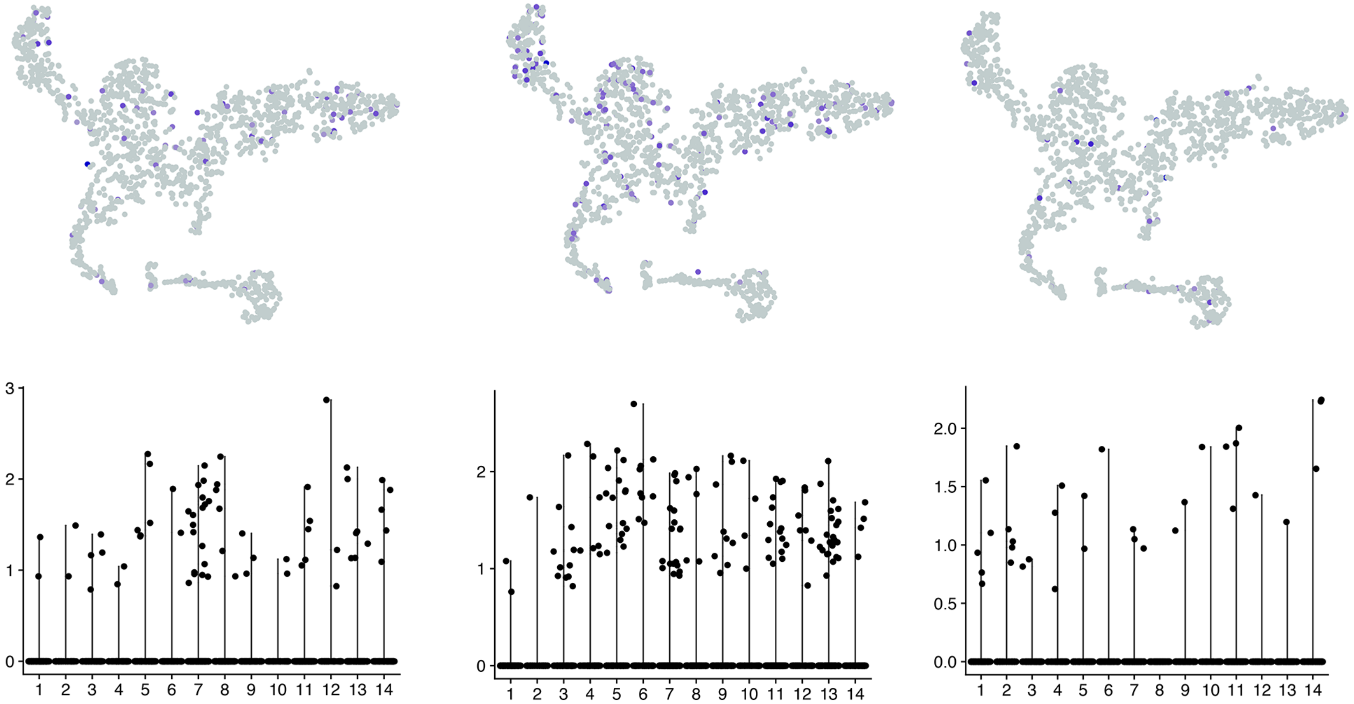

### Figure Supplementary 11 for Figure 6

Neuromasts derived from the Occipital prim and primD show a hair cell phenotype in *MZwnt11f1* mutants

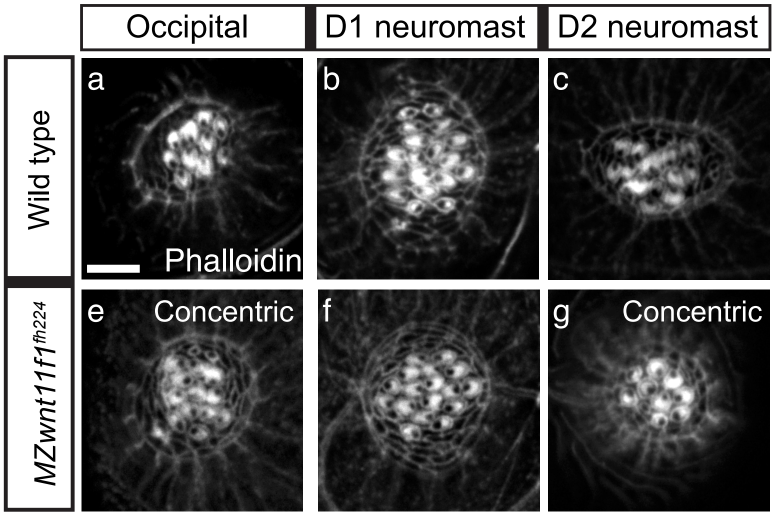

### Figure Supplementary 12

Distribution, but not ratio of expression, of Emx+ hair cells is affected in primII-derived neuromasts of *MZwnt11f1* mutants

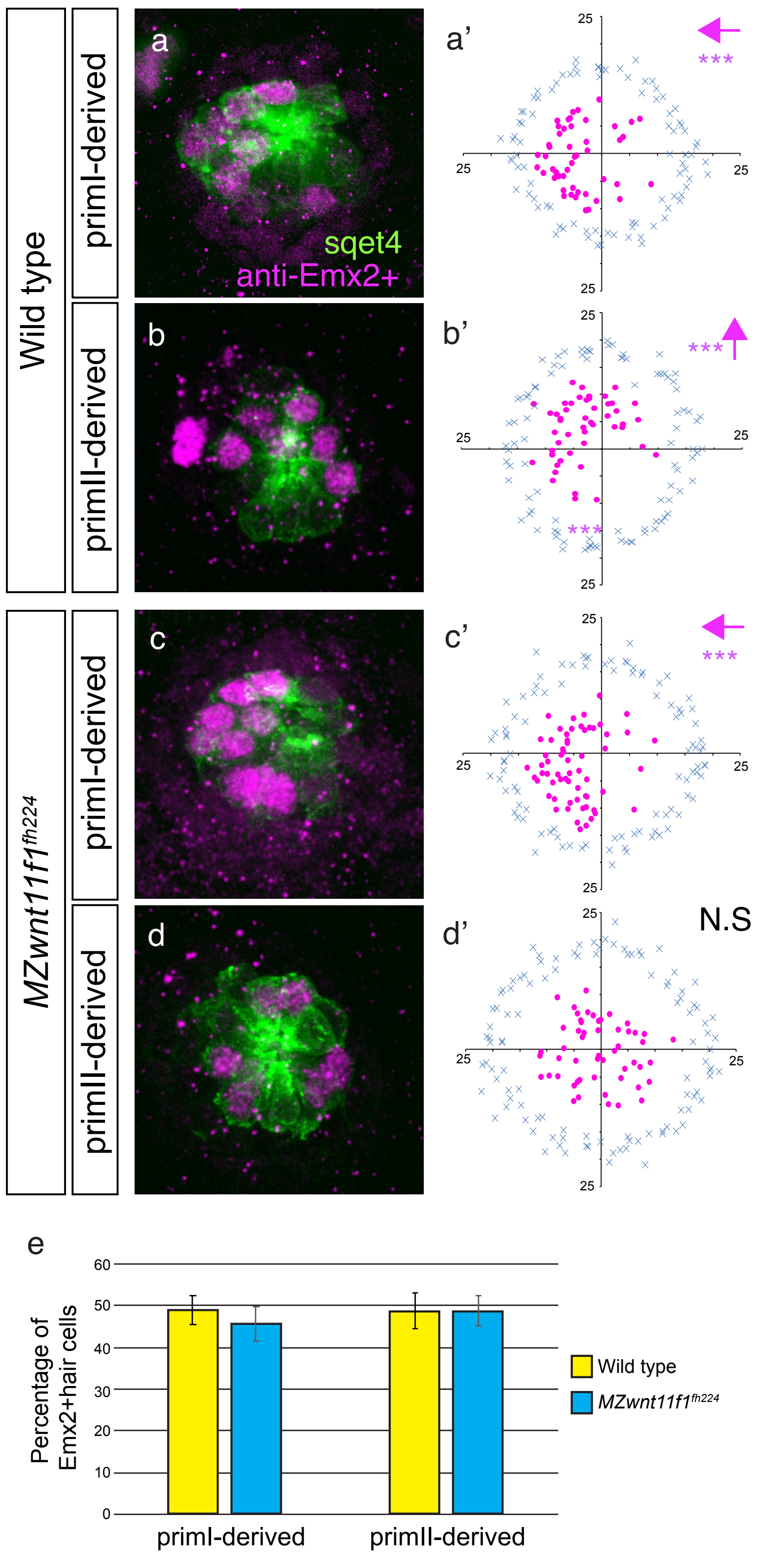
